# Targeting one-carbon metabolic vulnerabilities of metastasis with therapeutic potential

**DOI:** 10.64898/2026.02.07.704548

**Authors:** Yanlin Yu, Weiping Chen, Glenn Merlino

## Abstract

Evidence has shown that tumor progression is associated with the acquisition of growing autonomy and the creation of a complex signaling network through various signal pathways. Which particular signaling pathway is involved in the metastasis of a specific cancer is unclear. Here, we applied metastatic functional screening and identified that one-carbon and SSP metabolism pathways, as well as related genes, are associated with tumor metastasis inhibition. We engineered the cancer cells with poorly or highly metastatic potential to confirm the metabolism pathways regulating the ability to colonize different tissue sites. We also asked whether the restriction of the metabolism pathways by known inhibitors. We then identified three new compounds that can inhibit the expression of these genes and block tumor metastasis. Our findings uncovered a mechanism by which tumor cells reprogram their metabolism to promote migration, invasion, and survival at distant sites in tumor metastasis, offering a rational strategy to guide clinical treatment. The identified novel molecular proteins and pathways represent a promising therapeutic target for metastatic disease.

## Introduction

Metastasis is responsible for 90% of deaths in patients with cancer and is challenging to treat [1,2]. Available targeted therapies and immunotherapies have demonstrated promise for treating metastatic disease in some cancer patients [3,4]. However, the acquired resistance to targeted therapy, as well as a minority of patients displaying inefficiency in immunotherapies, hampers their durable efficacy [3,5,6]. Exploring the molecular mechanisms and vulnerable targets for treating metastasis remains the primary goal of current cancer research [3,7,8].

The metastatic process can be described as the outcome of interactions between tumor cells and the tumor microenvironment (TME), in which tumor cells undergo an evolutionary process to adapt and evade immunosurveillance [3,8–11]. Selective pressures of the tumor microenvironment and genetic alterations, such as oncogenic mutations and inactivation of tumor-suppressor genes, can select for tumor cells with the capability to grow despite the presence of environmental stressors, such as lack of oxygen and nutrients, low pH, reactive oxygen species, and the inflammatory response [11–13]. The acquisition of these functions can facilitate the initiation and progression of metastasis, where tumor cells develop local advantages in the primary tumor microenvironment and selective advantages during the adaptation and takeover of a distant organ microenvironment [8,11]. There is growing evidence that metastasis is a multidirectional process where tumor cells can seed at distant sites by multiple signaling pathways [8,9]. This behavior may explain the aberrant motility and invasiveness of tumor cells, the ability of stem-like cells to initiate tumors, and the interaction of stroma and tumors in metastasis [9,14]. Metastasis has also been likened to an ecological dispersal, where tumor cells disseminate, similar to diasporas in which people disperse from an established homeland [15,16], suggesting that a successful metastasis requires factors, such as an eligible primary tumor microenvironment, cancer cell migrants’ fitness, bidirectional ability to migrate between cancer sites, and suitable metastatic microenvironment sites. Metastases can also be described as ecological diseases [15–17]; foundational environmental principles, such as intraspecific (e.g., communication) and interspecific (e.g., competition, predation, parasitism, and mutualism) relationships, could help elucidate tumor progression and the context of tumor metastasis. The intraspecific and interspecific properties of metastasis are determined by the multiple networks of reprogramming.

The phenotypic complexity of the multistep cascade of metastasis formation contributes to the difficulties of identifying effective inhibitors against this deadly process. The success of cancer’s current treatment largely depends on the genetic mutations present within metastases. Genetically diverse metastases are more likely to harbor resistance mutations, contributing to treatment failure. Each metastasis is often assumed to be seeded precisely once, so its diversity cannot be attributed solely to genetic mutations. The shared vulnerability of the survival-required pathway would be the most common target for treating metastatic disease. Evidence shows that tumor progression is associated with increased autonomy and the formation of a complex signaling network through multiple signaling pathways. Tumor progression in different types of cancer involves widely variant signaling networks. Which particular signaling pathway is involved in the metastasis of a specific cancer is unclear. Identifying the precise signaling pathway or its molecular targets for mechanistic enlightenment would provide new strategies for clinical diagnosis and therapeutic utility, as well as prevent cancer metastasis. Here, we applied metastatic functional screening to discover metabolic vulnerabilities of metastatic cancer with therapeutic potential. We dissected this common pathway of metastases through the inhibition of metastasis phenotypes and identified the vulnerability of metabolism in metastasis for therapeutic utility.

## Results

### Metastasis treatment largely depends on genetic diversity

Due to the complex process of the metastatic cascade, treating metastatic disease is challenging because of the difficulty in identifying effective inhibitors against this deadly process. Here, we applied metastatic functional screening using molecular inhibitors in various metastatic cancers to identify targeted signaling pathways for combating metastasis. We first pretreated the metastatic cancer cells with different molecular inhibitors, including inhibitors of PKC (calphostin C, Cal C), Rho (Rho Inhibitor I, Rho), PKA (H-89), mTOR (Rapamycin, Rap), MAPK (PD98059, PD), PI3K/AKT (LY294002, LY), p38 (SB203580, SB) and HADC (TSA; MS-275, MS). Then, we examined the ability of metastasis in immunocompetent syngeneic or immunodeficent mouse models by tail vein injection (Figure 1A). Compared with the mock control, inhibition of PKC by Cal C or mTOR by rapamycin reduced the metastatic capability of all highly metastatic human A375sm and mouse B16F10 melanoma cell lines, as well as mouse rhabdomyosarcoma RMS14 (Figures 1B, 1C, and 1D), and human melanoma WM88, and mouse breast cancer 4T1 and rhabdomyosarcoma RMS10 (Figure S1). In contrast, inhibition of Rho, p38, and HADC did not affect the metastatic ability of these cells (Figures 1B, 1C, and 1D, S1). Interestingly, inhibition of MAPK only blocked the metastatic potential of RMS14 rhabdomyosarcoma cells, but not that of A375sm B16F10 melanoma cells (Figures 1B, 1C, and 1D). In contrast, the inhibition of PI3K/AKT and PKA only inhibited the metastatic potential of the melanoma cells, but not that of the rhabdomyosarcoma cell RMS14 (Figures 1B, 1C, and 1D). To further confirm the roles of PI3K/AKT and MAPK signaling in metastasis in melanoma and rhabdomyosarcoma, we pretreated the melanoma B16F10 cells with the PI3K/AKT inhibitor LY and the rhabdomyosarcoma RMS14 cells with the MAPK inhibitor PD at different times. Figures 1H and 1I showed that the metastatic potential gradually decreased following pretreatment from 12 to 48 hours. Although most inhibitors could inhibit cell proliferation, they did not match the phenotype of metastasis (Figures 1E, 1F, and 1G, S1), suggesting that the signaling pathways governing cell proliferation and metastasis are different. Our data demonstrated that PKC and mTOR inhibitors generally function to inhibit metastasis. Meanwhile, the function of MAPK and PI3K signaling pathways is cell type specific.

**Figure 1.**
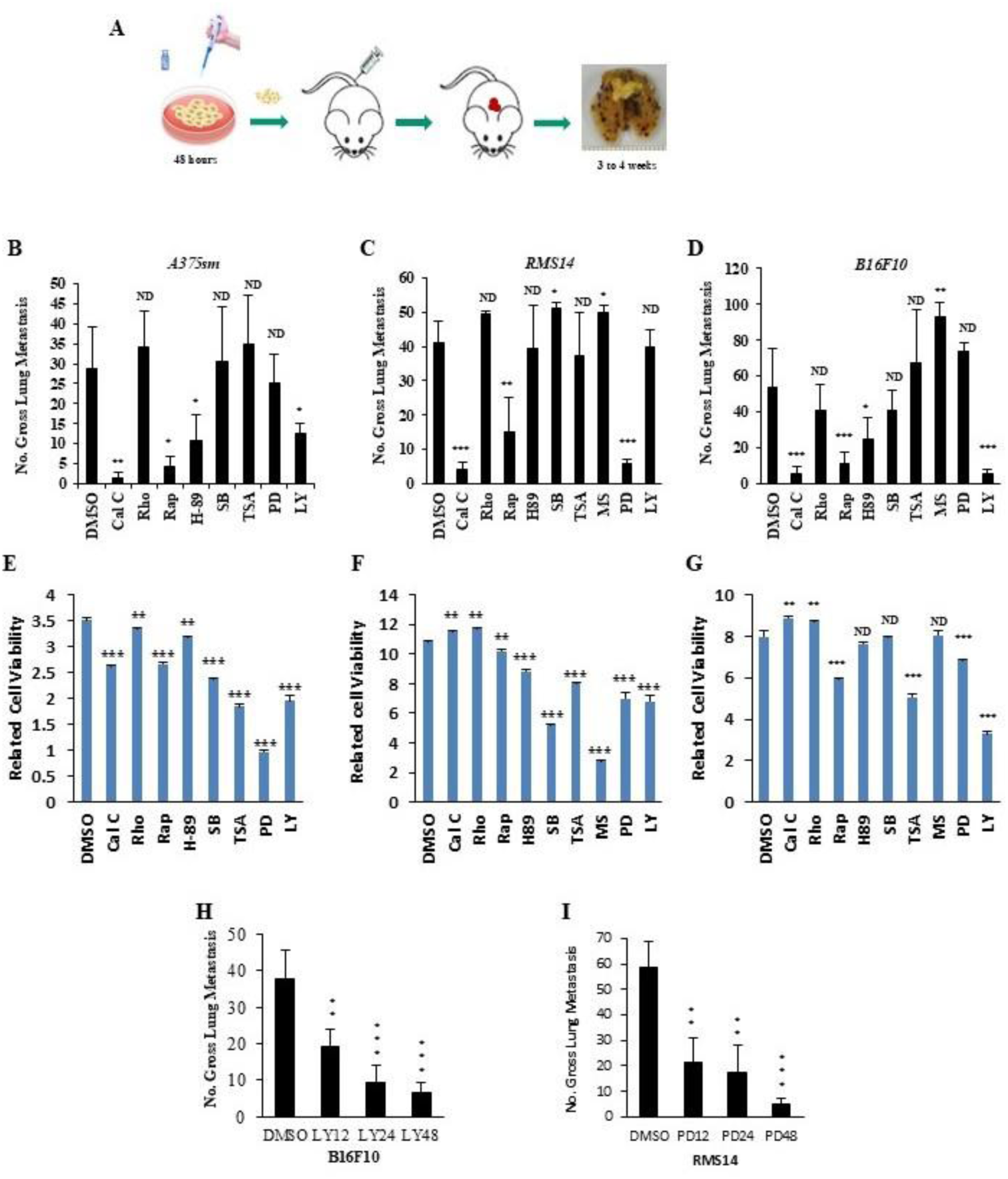
Metastasis intratumor and intertumor heterogeneity presented diverse responses to treatments. (**A**) Schematic of the metastasis functional screening using molecular inhibitors in vivo. (**B, C, and D**) Gross pulmonary metastases from human melanoma A375sm (B), mouse melanoma B16F10 (C), and mouse rhabdomyosarcoma RMS14 (D) in response to molecular inhibitor in the experimental metastasis assay by tail vein injection. Data are represented as mean ± SEM. The p-values were presented from an unpaired t-test analysis (two-tailed) compared with the control (DMSO). Cal C, 100 nM calphostin C; Rho, 12nM Rho Inhibitor I; Rap, 10nM Rapamycin; H-89, 50nM PKA inhibitor; SB, 10µM p38 inhibitor SB203580; TSA, 300nM HDAC inhibitor Trichostatin A; MS, 10µM HDAC inhibitor MS-275; PD, 20µM MAPK inhibitor PD98059; LY, 10µM PI3K/AKT inhibitor LY294002. *ND*, no statistical difference; \**p* < 0.05; \*\**p* < 0.01; \*\*\**p* < 0.001. n = 10. (**E, F, and G**) The related cell viability of human melanoma A375sm (E), mouse melanoma B16F10 (F), and mouse rhabdomyosarcoma (G) cells, pretreated with molecular inhibitors. Data are represented as mean ± SEM of three independent experiments. The p-values were presented from an unpaired t-test analysis (two-tailed) compared with the control (DMSO). *ND*, no statistical difference; \**p* < 0.05; \*\**p* < 0.01; \*\*\**p* < 0.001. (**H**) Gross pulmonary metastases from mouse melanoma B16F10 pretreated with 10µM PI3K/AKT inhibitor LY294002 at the indicated time (LY12, 12 hours, LY24, 24 hours and LY48, 48 hours). Data are represented as mean ± SEM. The p-values were presented from an unpaired t-test analysis (two-tailed) compared with the control (DMSO). *ND*, no statistical difference; \**p* < 0.05; \*\**p* < 0.01; \*\*\**p* < 0.001. n=10. (**I**) Gross pulmonary metastases from mouse rhabdomyosarcoma RMS14 cells pretreated with 20µM MAPK inhibitor PD98059 at the indicated time (PD12, 12 hours, PD24, 24 hours and PD48, 48 hours). Data are represented as mean ± SEM. The p-values were presented from an unpaired t-test analysis (two-tailed) compared with the control (DMSO). *ND*, no statistical difference; \**p* < 0.05; \*\**p* < 0.01; \*\*\**p* < 0.001. n=10.

### Identification of metabolism pathways in metastases

The diversity of the signaling pathways supporting cancer metastasis limits the possibility of designing overarching treatment strategies. The common signaling pathways underlying the phenotype of metastatic inhibition in various cancers may provide information for designing a new approach to combating metastatic tumors. To focus on the inhibition of tumor metastasis, we next used microarray analysis to compare the metastasis inhibition groups with the control group in both melanoma B16F10 (treated with control, Cal A, Rap, LY24, and LY 48) and rhabdomyosarcoma RMS14 (treated with control, Cal A, Rap, PD24, and PD 48) (Figure 2A). We identified 46 genes that were significantly differentially expressed in both types of cancer cell groups (Figure 2B). Twenty-four of them were upregulated considerably, including tumor suppressors such as Ypel3 [18], HDAC11, and Bim, while 25 genes were remarkably downregulated, including oncogenes or metastasis enhancers such as Ezrin [19], Zwint, and Axl (Figure 2B). Interestingly, except for oncogene or metastasis enhancers in 25 downregulated genes, 12 downregulated genes are involved in metabolism signaling pathways and survival, including Glut3, LRP8, MCT1, Enoph1, PHGDH, Nqo1, PSAT1, SHMT2, MTHFD2, MTHFD1, PSPH, and BCAT1 (Figure 2B). Through gene functional pathway analysis, we identified the top 15 significantly changed functional pathways that involve the serine, cysteine, and glycine family amino acid metabolic process, as well as the methionine biosynthetic process, which were significantly boosted in all metastatic tumors (Figure 2C). Gene signaling pathway analysis showed that these genes regulate the same one-carbon metabolism and De novo serine synthesis (SSP) (Figure 2D).

**Figure 2.**
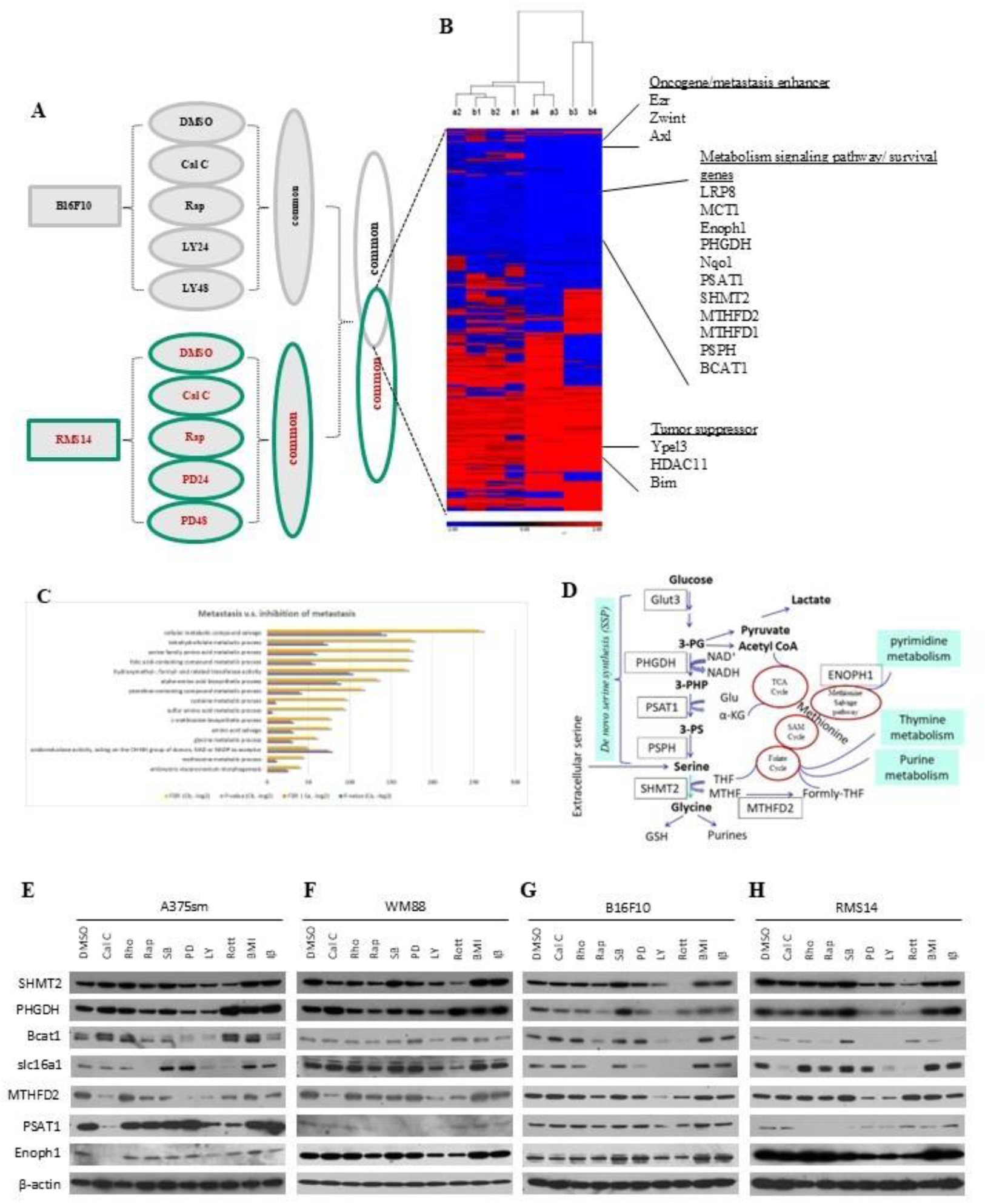
Identification of metabolism pathways in metastases. (A) Schematic of microarray analysis to focus on the inhibition of tumor metastasis. (B) The significant differentiated gene expression patterns associated with the inhibition of tumor metastasis progression and the identification of metabolism pathways associated with tumor metastasis. The significantly expressed genes in B16F10 (a) or RMS14 (b) cells treated with inhibitors (Cal C, 100 nM calphostin; Rap, 10nM Rapamycin; 20µM MAPK inhibitor PD98059; LY, 10µM PI3K/AKT inhibitor LY294002) compared with DMSO control were filtered by a p-value of 0.05 and an absolute value of fold change of 1.5 in ANOVA analysis. a1, Cal C v.s. DMSO; a2, Rap v.s. DMSO; a3, LY24 v.s. DMSO; a4, LY48 v.s. DMSO in B16F10 cells. b1, Cal C v.s. DMSO; b2, Rap v.s. DMSO; b3, PD24 v.s. DMSO; b4, PD48 v.s. DMSO in RMS14 cells. (**C**) Gene signaling pathway (GO gene pathway) analysis revealed that top 15 biological pathways and functions within a given gene list from micrroarry analysis regulate one-carbon metabolism and de novo serine synthesis (SSP). Both *p* and FDR valuest were transformed to negative log (-log2) values. (**D**) Schematic of the one-carbon metabolism and De novo serine synthesis (SSP) pathways. (**E to H**) Validation of gene expression identified in cDNA microarray analysis by Western blot analysis. The human melanoma A375sm (E) and WM88 (F), mouse melanoma B16F10 (G), or mouse rhabdomyosarcoma RMS14 (H) cells were treated with molecular inhibitors. DMSO, as a mock control; Cal C, 100 nM calphostin C; Rho, 12nM Rho Inhibitor I; Rap, 10nM Rapamycin; SB, 10 µM p38 inhibitor SB203580; PD, 20 µM MAPK inhibitor PD98059; LY, 10 µM PI3K/AKT inhibitor LY294002; Rott, 5μM of rottlerin; BMI, 20 nM of bisindoylmaleimide I; Iβ, 20 nM of PKCβ inhibitor I. *β*-actin was used as a control.

To validate the microarray analysis findings, we treated the four highly metastatic human and mouse cells with various inhibitors. Then, we examined the protein levels of the genes in the one-carbon or SSP metabolism signaling pathway. As Figures 2E to 2H show, treatment of Cal C, Rott, and Rap could decrease the expression of one-carbon metabolism genes in all human (A375sm and WM88) and mouse (B16F10, RMS14) highly metastatic tumor cell lines. At the same time, LY inhibitor downregulated the one-carbon metabolism genes expression in melanoma (Figure 2E, 2F, and 2G), and PD reduced the one-carbon metabolism genes expression in rhabdomyosarcoma cells (Figure 2H). Our data confirmed the finding from our microarray analysis, suggesting that one-carbon metabolism genes regulated by these inhibitors may be related to metastatic inhibition in both melanoma and rhabdomyosarcoma.

### One carbon metabolism-related gene is associated with the potential for metastasis

ENOPH1 is an essential enzyme in the methionine salvage pathway, which maintains methionine levels in vivo by recycling the one-carbon unit and mitigating stress [20]. Therefore, it regulates the rate of protein synthesis (Figure S2A). To further confirm the function of one-carbon metabolism in metastasis, we studied the function of ENOPH1 in regulating tumor metastasis using different strategies (Figure 3A). We first overexpressed ENOPH1 in poorly metastatic melanoma B16F1 cells and then tested its metastatic potential in syngeneic immunocompetent mice by tail vein injection (Figure 3B). As shown in Figure 3C, stable overexpression of ENOPH1 in poorly metastatic melanoma B16F1 cells promotes metastasis (Figure 3C) and cell proliferation (Figure 3D). In contrast, the knockdown of ENOPH1 by shRNA for ENOPH1 in highly metastatic melanoma B16F10 cells reduced the metastatic potential (Figure 3E, 3F) and cell proliferation (Figure 3G). To further confirm that downregulated ENOPH1 inhibits tumor metastasis, we depleted ENOPH1 using TALEN and CRISPR/Cas9 gene editing technologies (Figures 3H, 3K). Then, we corroborated that the deficiency of ENOPH1 in highly metastatic melanoma B16F10 cells significantly reduced lung metastasis (Figures 3I and 3L) and cell proliferation (Figures 3J and 3M). Using metabolon analysis, we noted that overexpression of ENOPH1 promoted the TCA cycle, serine, glycine, methionine, and folate metabolism; in contrast, the knockout of ENOPH1 downregulated the TCA cycle, serine, glycine, methionine, and folate metabolism (Figures 3N, 3O, and 3P). Seahorse analysis confirmed that overexpression of ENOPH1 enhances both basal and compensatory glycolysis, ATP production, and OCR (Figures 4A-D), whereas knockdown or knockout inhibits glycolysis, ATP production, and OCR (Figures 4E-L).

**Figure 3.**
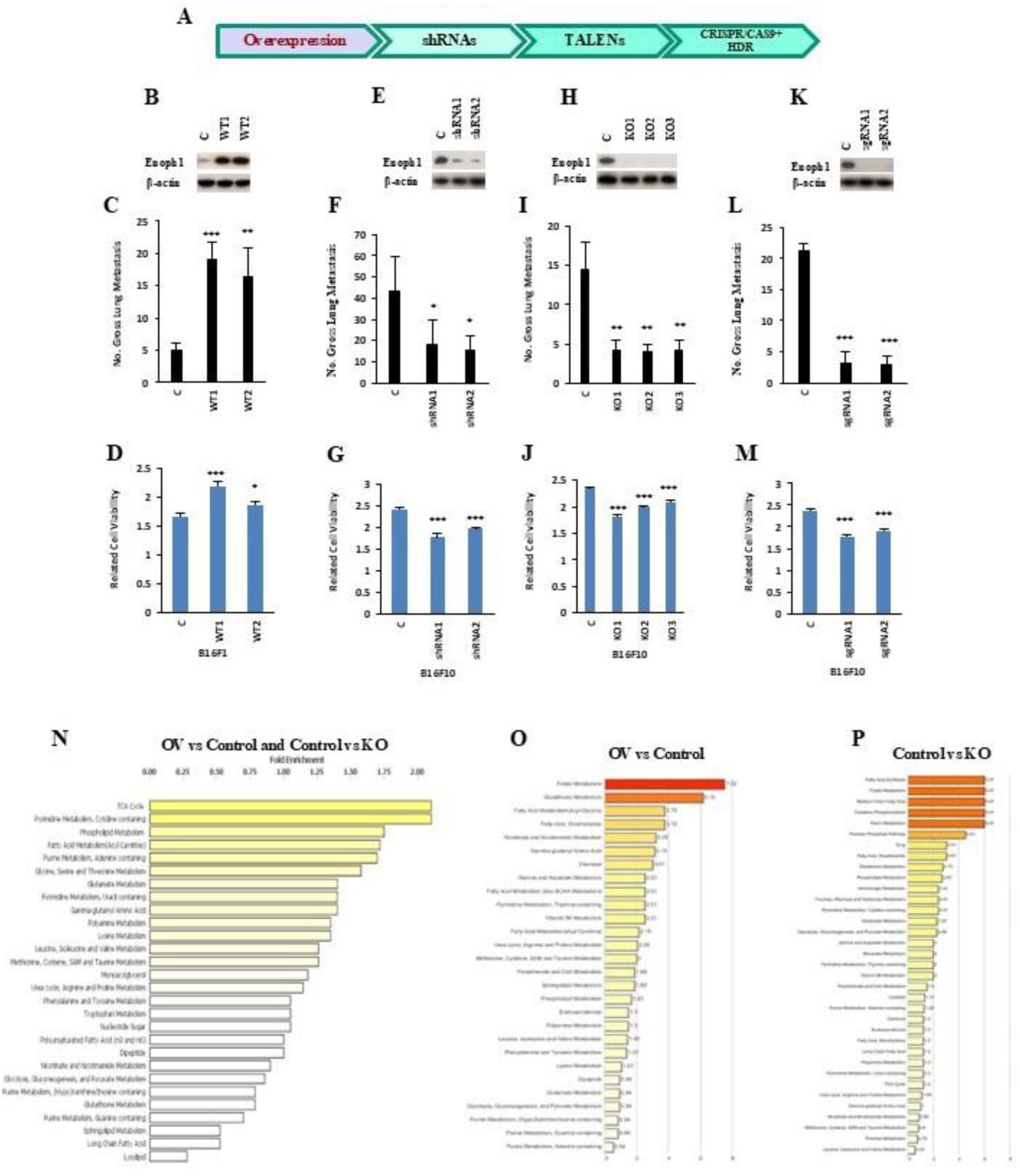
One carbon metabolism-related gene ENOPH1 is associated with the potential for metastasis. (**A**) The schematic procedure of ENOPH1 functional studies. (**B**) Western blotting showed overexpression of ENOPH1 in poorly metastatic mouse melanoma B16F1 cells. Protein levels of ENOPH1 in B16F1 cells transfected with the wildtype ENOPH1 gene (WT1 and WT2). C, empty vector; WT1 and WT2, two different clones with ENOPH1 wildtype. The protein level of *β*-actin is the loading control. (**C**) Gross pulmonary metastases from mouse melanoma B16F1 cells transfected with the wildtype of the ENOPH1 gene (WT1 and WT2) by tail vein injection. C, empty vector; WT1 and WT2, two different clones with ENOPH1 wildtype. (**D**) Effects on cell growth by ENOPH1 overexpression. The data represent three independent experiments. Graphs show the mean ± SEM. The p-value is shown by an unpaired t-test (two-tailed). c, empty vector; WT1 and WT2, ENOPH1 expression plasmid. *ND*, no statistical difference; \**p* < 0.05; \*\**p* < 0.01; \*\*\**p* < 0.001. (**E**) Western blotting showed the knockdown of ENOPH1 in highly metastatic mouse melanoma B16F10 cells using two shRNAs targeting ENOPH1. Protein levels of ENOPH1in B16F10 cells transfected with shRNAs ENOPH1 plasmids (shRNA1 and shRNA2). C, empty vector; shRNA1 and shRNA2, two different shRNAs for ENOPH1. The protein level of *β*-actin is the loading control. (**F**) Gross pulmonary metastases from mouse melanoma B16F10 cells transfected with shRNA ENOPH1 plasmids (shRNA1 and shRNA2) by tail vein injection. C, empty vector; shRNA1 and shRNA2, two different ENOPH1 shRNAs. (**G**) Effects on cell growth by shRNA ENOPH1 knockdown. The data represent three independent experiments. Graphs show the mean ± SEM. The p-value is shown by an unpaired t-test (two-tailed). c, empty vector; shRNA1 and shRNA2 for ENOPH1 expression plasmids. *ND*, no statistical difference; \**p* < 0.05; \*\**p* < 0.01; \*\*\**p* < 0.001. (**H**) Western blotting showed the knockout of ENOPH1 in highly metastatic mouse melanoma B16F10 cells using Transcription Activator-Like Effector Nucleases (TALENs) genomic editing technology for ENOPH1. Protein levels of ENOPH1 in B16F10 cells transfected with TALEN knockout plasmids for ENOPH1 (KO1, KO2, and KO3). C, empty vector; KO1, KO2, and KO3, three different clones with TALEN knockout plasmid for ENOPH1. The protein level of *β*-actin is the loading control. (**I**) Gross pulmonary metastases from mouse melanoma B16F10 cells transfected with TALEN ENOPH1 plasmids (KO1, KO2, and KO3) by tail vein injection. C, empty vector; KO1, KO2, and KO3, three different clones with TALEN knockout plasmid for ENOPH1. (**J**) Effects on cell growth by TALENs knockout for ENOPH1. The data represent three independent experiments. Graphs show the mean ± SEM. The p-value is shown by an unpaired t-test (two-tailed). c, empty vector; KO1, KO2 and KO3, three different clones with TALEN knockout plasmid for ENOPH1. (**K**) Western blotting showed the knockout of ENOPH1 in highly metastatic mouse melanoma B16F10 cells using CRISPR/Cas9 gene editing technology for ENOPH1. Protein levels of ENOPH1 in B16F10 cells transfected with CRISPR/CAS9 knockout plasmids for human ENOPH1 (sgRNA1 and sgRNA2). C, empty vector; sgRNA1 and sgRNA2, two different CRISPR/CAS9 knockout plasmids for ENOPH1. The protein level of *β*-actin is the loading control. (**L**) Gross pulmonary metastases from mouse melanoma B16F10 cells transfected with CRISPR/CAS9 ENOPH1 plasmids (sgRNA1 and sgRNA2) by tail vein injection. C, empty vector; sgRNA1 and sgRNA2, two different CRISPR/CAS9 knockout plasmids for ENOPH1. (**M**) Effects on cell growth by CRISPR/CAS9 knockout for ENOPH1. The data represent three independent experiments. Graphs show the mean ± SEM. The p-value is shown by an unpaired t-test (two-tailed). c, empty vector; sgRNA1 and sgRNA2, two different CRISPR/CAS9 knockout plasmids for ENOPH1. *ND*, no statistical difference; \**p* < 0.05; \*\**p* < 0.01; \*\*\**p* < 0.001. (**N to P**) The metaboton analysis interrogates biochemical profiles in overexpression or knockout of ENOPH1 tumor cells (The Characterizing metabolism associated with altered enolase expression). (N) Global biochemical profiles of the combination ENOPH1 overexpression v.s. control in B16F1 cells (OV vs Control) and ENOPH1 control v.s. Knockout in B16F10 cells (Control vs KO). (O) The biochemical metabolite profiles of overexpression v.s. control in B16F1 cells (OV vs Control). (P) The biochemical metabolite profiles of ENOPH1 control v.s. knockout in B16F10 cells (Control vs KO). Using metabolon analysis, we observed that overexpression of ENOPH1 promoted the TCA cycle, as well as serine, glycine, methionine, and folate metabolism. In contrast, the knockout of ENOPH1 downregulated these metabolic pathways.

**Figure 4.**
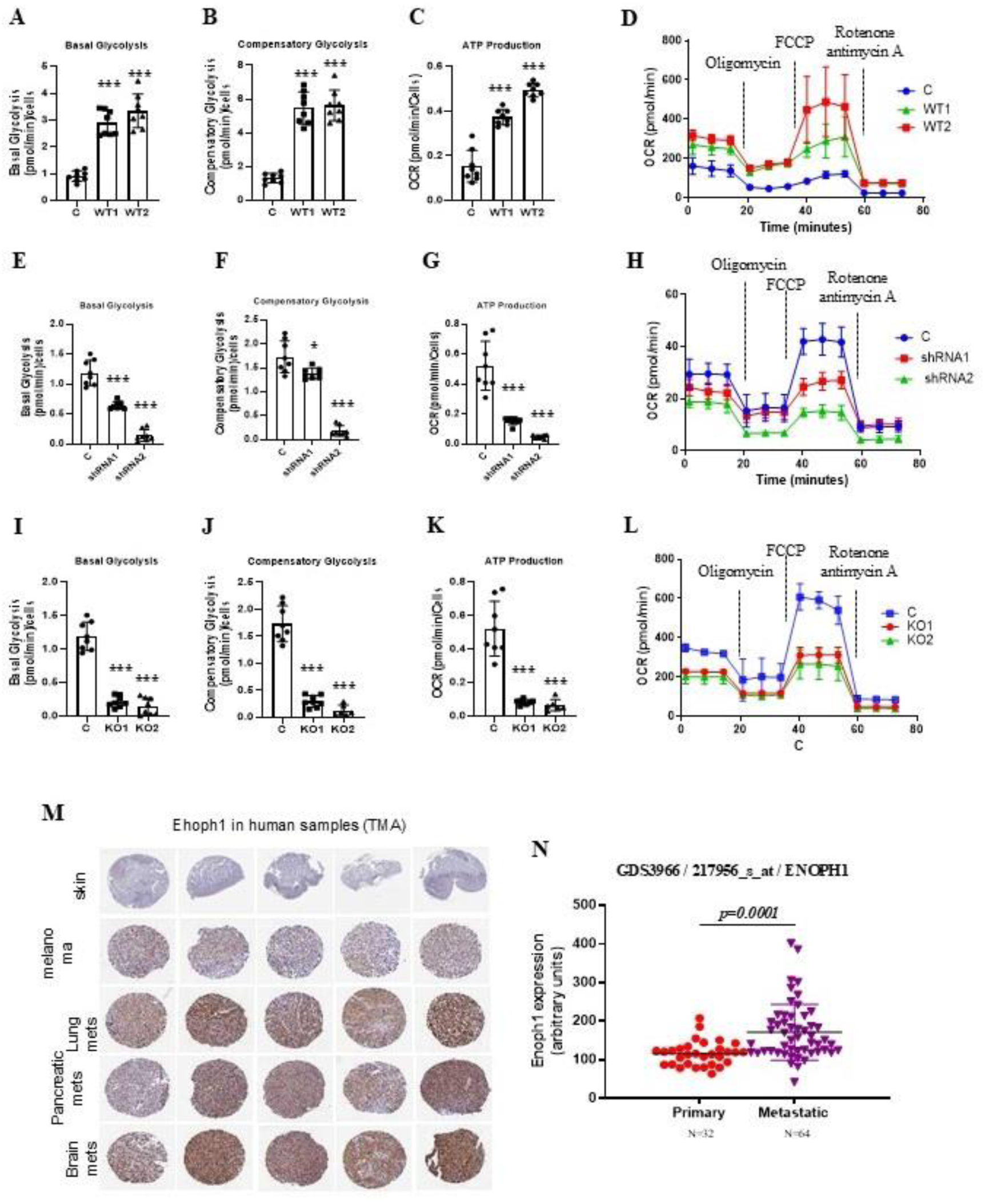
The tumor metabolism and metastasis are associated with altered enolase expression. (**A, B, C, D**) Seahorse analysis confirmed that overexpression of ENOPH1 in B16F1 cells enhances both basal (A) and compensatory glycolysis (B), ATP production (C), and OCR (D). C, empty vector control; OV1 and OV2, the cells expressing the ENOPH1 expression plasmid. The data represent three independent experiments. Graphs show the mean ± SEM. The p-value is shown by an unpaired t-test (two-tailed). *ND*, no statistical difference; \**p* < 0.05; \*\**p* < 0.01; \*\*\**p* < 0.001. (**E, F, G, H**) Seahorse analysis confirmed that knockdown of ENOPH1 in B16F10 cells reduces both basal (E) and compensatory glycolysis (F), ATP production (G), and OCR (H). C, empty vector control; shRNA1 and shRNA2, the cells expressing the ENOPH1 shRNA plasmids. The data represent three independent experiments. Graphs show the mean ± SEM. The p-value is shown by an unpaired t-test (two-tailed). *ND*, no statistical difference; \**p* < 0.05; \*\**p* < 0.01; \*\*\**p* < 0.001. (**I, J, K, L**) Seahorse analysis confirmed that knockout of ENOPH1 in B16F10 cells reduces both basal (I) and compensatory glycolysis (J), ATP production (K), and OCR (L). C, empty vector control; KO1 and KO2, the cells expressing the ENOPH1 TALEN-deficient plasmids. The data represent three independent experiments. Graphs show the mean ± SEM. The p-value is shown by an unpaired t-test (two-tailed). *ND*, no statistical difference; \**p* < 0.05; \*\**p* < 0.01; \*\*\**p* < 0.001. (**M**) ENOPH1 is significantly expressed in metastatic melanoma, including lung, pancreatic, and brain metastasis, compared with normal skin and melanoma by tissue microarray (TMA) analysis. (**N**) ENOPH1 is highly significantly expressed in metastatic melanoma compared to primary melanoma in the GDS3966 dataset. Graphs show the mean ± SEM. The p-value is shown by an unpaired t-test (two-tailed).

We found that ENOPH1 is significantly expressed in metastatic melanoma, including lung, pancreatic, and brain metastasis, compared with normal skin and melanoma by tissue microarray (TMA) analysis (Figure 4M). Additionally, ENOPH1 is highly significantly expressed in metastatic melanoma compared to primary melanoma in the GDS3966 dataset (Figure 4N). Moreover, the high level of ENOPH1 expression is associated with worse patient survival in various types of tumors in the TCGA datasets (Figure S2). Our data demonstrates that a high level of ENOPH1 is associated with metastatic potential, and inhibition of ENOPH1 expression could reduce metastasis, suggesting that ENOPH1 can be a target for treating metastatic diseases.

### De novo serine synthesis (SSP) metabolism-related genes associate with metastatic potential

We also found that downregulated genes in the SSP metabolism signaling pathway are associated with metastasis inhibition (Figures 2B and 2D). Among them, PHGDH (phosphoglycerate dehydrogenase) is the first rate-limiting enzyme in the pathway (Figure 2D) and is amplified in many cancers, including melanoma and breast cancer [21–23]. To confirm the function of the SSP metabolism signaling pathway in metastasis, we investigated the role of PHGDH in metastasis. We first introduced the PHGDH gene into poorly metastatic human melanoma A375p cells (Figure 5A). Then, we tested its metastatic potential in two stable overexpressing PHGDH clones and an empty vector control in NSG mice by tail vein injection and orthotopic footpad injection. As Figures 5B and 5C show, overexpression of PHGDH can significantly enhance lung and liver metastasis using either tail vein injection (Figure 5B) or orthotopic footpad injection (Figure 5C). To better track tumor metastases, we transfected wild type and a dominant active form (DA) of PHGDH into Akluc [24] labeled A375p cells (Figure 5D). As expected, expressing both wild-type and dominant active forms significantly promoted metastasis of A375p cells in the lung (Figure 5E) and intensity of lung image as luminescence by In Vivo Imaging System (IVIS) through the tail vein and orthotopic footpad injection (Figures 5F). Also, overexpression of PHGDH in the poorly metastatic mouse melanoma B16F1 was found to increase lung metastasis in C57BL/6 syngeneic, immunocompetent mice (Figure 5G).

**Figure 5.**
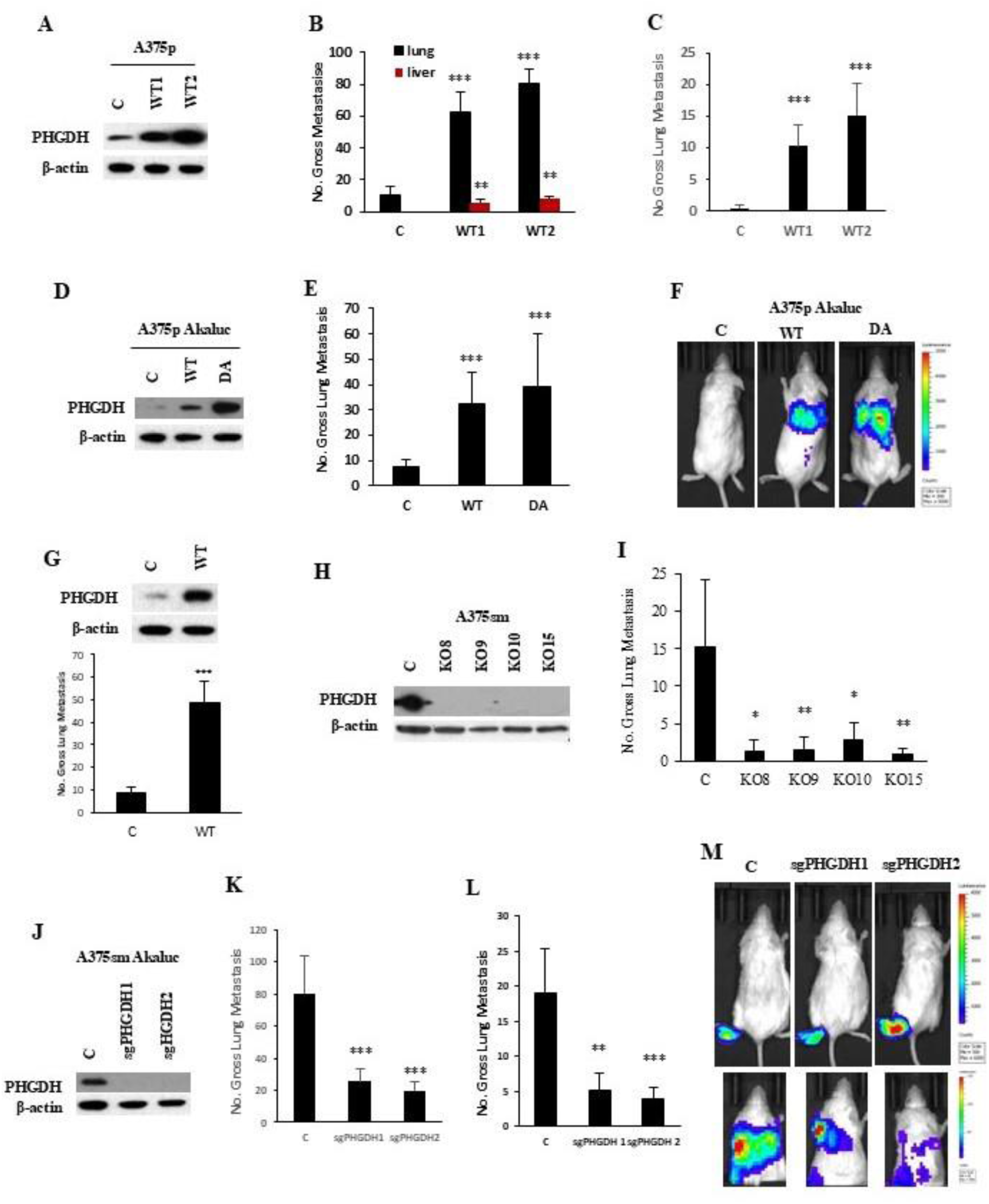
De novo serine synthesis (SSP)-related metabolism genes are associated with the potential for metastasis. (**A**) Western blotting showed overexpression of PHGDH (a rate-limiting enzyme in the SSP pathway) in poorly metastatic human melanoma A375p cells. Protein levels of PHGDH in A375p cells transfected with the wildtype PHGDH gene (WT1 and WT2). C, empty vector; WT1 and WT2, two different clones with PHGDH wildtype. The protein level of β-actin is the loading control. (**B**) Gross pulmonary metastases from human melanoma A375p cells transfected with the wildtype PHGDH gene (WT1 and WT2) were induced by tail vein injection. C, empty vector; WT1 and WT2, two different clones with PHGDH wildtype. N=10, Graphs show the mean ± SEM. The p-value is shown by an unpaired t-test (two-tailed). *ND*, no statistical difference; \**p* < 0.05; \*\**p* < 0.01; \*\*\**p* < 0.001. (**C**)) Gross pulmonary metastases from human melanoma A375p cells transfected with wildtype PHGDH gene (WT1 and WT2) by orthotopic footpad injection. C, empty vector; WT1 and WT2, two different clones with PHGDH wildtype. N =10. Graphs show the mean ± SEM. The p-value is shown by an unpaired t-test (two-tailed). *ND*, no statistical difference; \**p* < 0.05; \*\**p* < 0.01; \*\*\**p* < 0.001. (**D**) Western blotting showed overexpression of PHGDH wildtype (WT) and dominant active form (DA) in poorly metastatic human melanoma A375p cells with Akaluc (A375p-Akaluc). Protein levels of PHGDH in A375p-Akaluc cells transfected with PHGDH wildtype (WT) or dominant active form (DA). C, empty vector. The protein level of β-actin is the loading control. (**E**) Gross pulmonary metastases from human melanoma A375p-Akaluc cells transfected with PHGDH wildtype (WT) or dominant active form (DA) gene by tail vein injection. C, empty vector. N=10. Graphs show the mean ± SEM. The p-value is shown by an unpaired t-test (two-tailed). *ND*, no statistical difference; \**p* < 0.05; \*\**p* < 0.01; \*\*\**p* < 0.001. (**F**) Representation of lung bioluminescence and fluorescence images in mice with A375p-Akaluc cells transfected with vector control (c) or wildtype PHGDH (WT) and dominant active PHGDH (DA) by In Vivo Imaging System (IVIS). (**G**) Top panel: Western blotting showed overexpression of PHGDH wildtype (WT) or empty vector control (c) in poorly metastatic mouse melanoma B16F1 cells. Bottom panel: Gross pulmonary metastases from mouse melanoma B16F1 transfected with wildtype PHGDH (WT) or empty vector control (c). N=10, Graphs show the mean ± SEM. The p-value is shown by an unpaired t-test (two-tailed). *ND*, no statistical difference; \**p* < 0.05; \*\**p* < 0.01; \*\*\**p* < 0.001. (**H**) Western blotting showed deficient PHGDH expression by CRISPR/CAS9 gene editing in highly metastatic human melanoma A375sm cells. C, empty vector control; KO8, KO9, KO10 and KO15, different knockout clones. β-actin as a protein internal loading control. (**I**) Gross pulmonary metastases from human melanoma A375sm cells transfected with CRISPR/CAS9 knockout PHGDH plasmids in NSG mice by orthotopic footpad injection. C, empty vector; KO8, KO9, KO10 and KO15, different PHGDH knockout clones. N=10, Graphs show the mean ± SEM. The p-value is shown by an unpaired t-test (two-tailed). *ND*, no statistical difference; \**p* < 0.05; \*\**p* < 0.01; \*\*\**p* < 0.001. (**J**) Western blotting showed the knockout of PHGDH by CRISPR/CAS9 sgRNA gene editing in highly metastatic human melanoma A375sm cells labeled with Akaluc (ref) (A375sm-Akaluc). Protein levels of PHGDH in A375sm-Akaluc cells transfected with CRISPR/CAS9 plasmids (sgRNA1 and sgRNA2). C, empty vector. The protein level of β-actin is the loading control. (**K**) Gross pulmonary metastases from human melanoma A375sm-Akaluc cells transfected with PHGDH sgRNA (sgRNA1 and sgRNA2) in NSG mice by tail vein injection. C, empty vector. N=10, Graphs show the mean ± SEM. The p-value is shown by an unpaired t-test (two-tailed). *ND*, no statistical difference; \**p* < 0.05; \*\**p* < 0.01; \*\*\**p* < 0.001. (**L**) Gross pulmonary metastases from human melanoma A375sm-Akaluc cells transfected with PHGDH sgRNA (sgRNA1 and sgRNA2) in NSG mice by orthotopic footpad injection. C, empty vector. N=10, Graphs show the mean ± SEM. The p-value is shown by an unpaired t-test (two-tailed). *ND*, no statistical difference; \**p* < 0.05; \*\**p* < 0.01; \*\*\**p* < 0.001. (**M**) Representation of lung bioluminescence and fluorescence images in mice with A375p-Akaluc cells transfected with either a vector control (c) or CRISPR/CAS9 sgRNA for PHGDH (sgRNA1 and sgRNA2) using an In Vivo Imaging System (IVIS). C, empty vector control.

To confirm our findings, we also knocked out PHGDH in highly metastatic human melanoma A375sm cells using CRISPR/Cas9 technology (Figure 5H). When the PHGDH-deficient cells were injected into NSG mice by orthotopic footpad injection or tail vein injection, the PHGDH-deficient cells showed a significant reduction in metastasis in NSG mice (Figure 5I). To be able to track the tumor metastasis and confirm the findings, we also knocked out PHGDH in highly metastatic human melanoma A375sm with Akaluc (A375smAkaluc) cells using CRISPR/Cas9 technology (Figure J). The knockout of PHGDH significantly reduced experimental and orthotopic lung metastasis by tail vein injection and orthotopic footpad injection (Figures 5K and 5L), as well as decreased the counts of luminescence in the lung by In Vivo Imaging System (IVIS) (Figure 5M). Additionally, overexpression of PHGDH promoted glycolysis, ATP production, and OCR activation (Figures 6A-D), whereas knockout of PHGDH inhibited glycolysis, ATP production, and OCR activation (Figures 6E-H). We next labeled the A375sm cells with Akaluc and tracked the tumor cells with PHGDH deficiency in vivo after tumor cell transplantation. We found that all cells reached the lung with similar luminescence intensity after cell injection within 30 minutes (Figure 6I). However, in one week, the control group with A375sm cells still maintained a similar luminescence intensity as early as 30 minutes, while the group with PHGDH-deficient A375sm cells lost the luminescence intensity dramatically (Figure 6J), implicating that loss of PHGDH affects the tumor cell survival in the lung, and then reduces the lung metastasis (Figure 6K).

**Figure 6.**
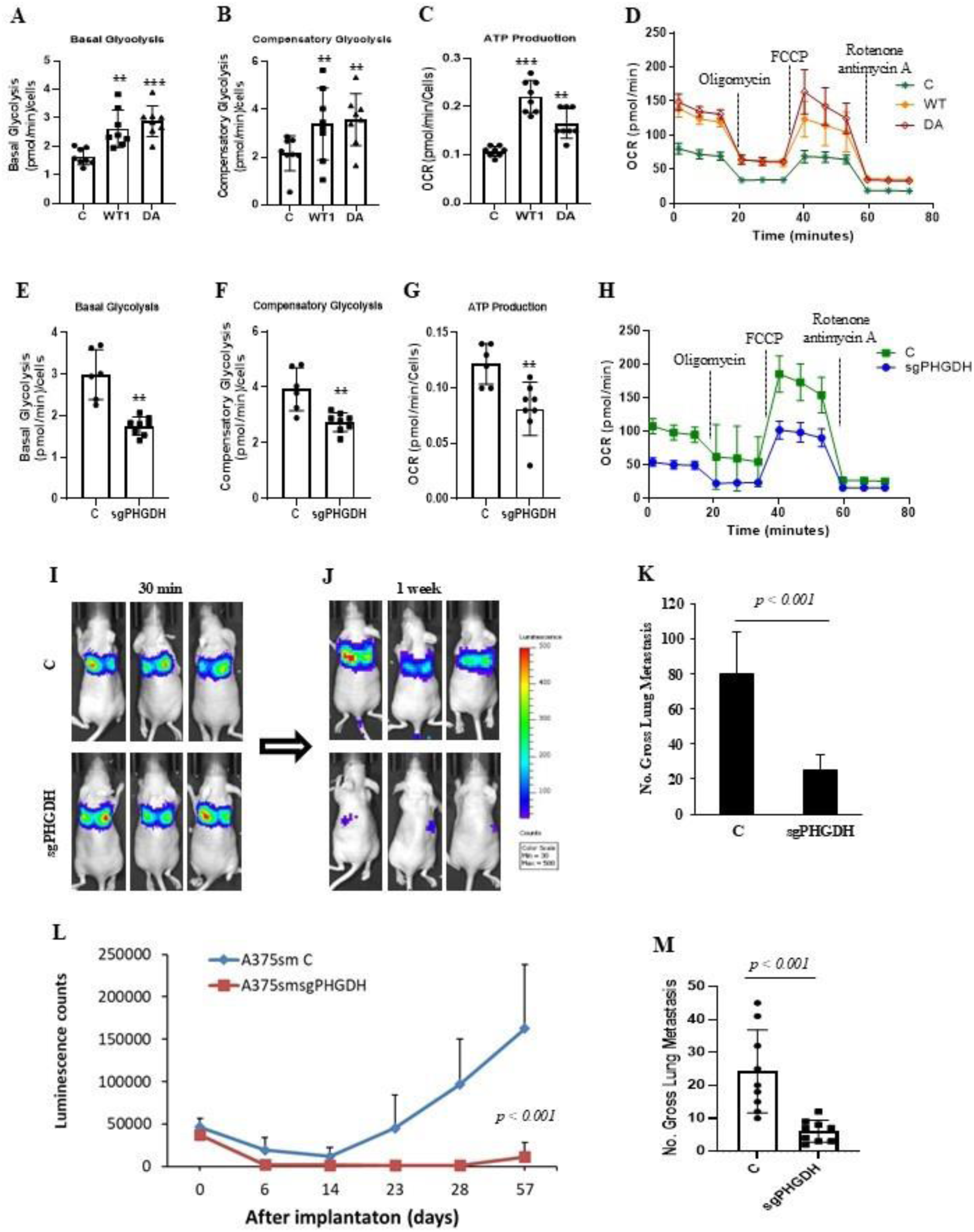
The alteration of PHGDH expression affects the tumor cell metabolism, survival in vivo, as well as the potential of metastasis. (**A, B, C, D**) Seahorse analysis confirmed that overexpression of PHGDH wildtype (WT) or dominant active form (DA) in A375p cells enhances both basal (A) and compensatory glycolysis (B), ATP production (C), and OCR (D). C, empty vector control. The data represent three independent experiments. Graphs show the mean ± SEM. The p-value is shown by an unpaired t-test (two-tailed). *ND*, no statistical difference; \**p* < 0.05; \*\**p* < 0.01; \*\*\**p* < 0.001. (**E, F, G, H**) Seahorse analysis confirmed that PHGDH deficiency (sgRNA) by CRISPR/CAS9 sgRNA in A375sm cells reduces both basal (E) and compensatory glycolysis (F), ATP production (G), and OCR (H). C, empty vector control. The data represent three independent experiments. Graphs show the mean ± SEM. The p-value is shown by an unpaired t-test (two-tailed). *ND*, no statistical difference; \**p* < 0.05; \*\**p* < 0.01; \*\*\**p* < 0.001. (**I, J**) Representation of lung bioluminescence and fluorescence images to track A375sm-Akaluc tumor cells transfected with vector control (C) or CRISPR/CAS9 sgRNA for PHGDH (sgRNA) at 30 min (I) and 1 week (J) after tail vein injection by In Vivo Imaging System (IVIS). (**K**) Gross pulmonary metastases from human melanoma A375sm-Akaluc cells transfected with PHGDH sgRNA (sgRNA) in NSG mice by tail vein injection. C, empty vector. N=10. Graphs show the mean ± SEM. The p-value is shown by an unpaired t-test (two-tailed). (**L**) The data showed the bioluminescence signals of tumor metastasis in NSG mice, tracked by the In Vivo Imaging System (IVIS), for 57 days after transplantation. C, the A375sm-Akaluc cells transfected with the control plasmid; sgRNA, the A375sm-Akaluc cells transfected with the CRISPR/CAS9 sgRNA for the PHGDH plasmid. Ten mice per group (N=10). Graphs show the mean ± SEM. The p-value is shown by an unpaired t-test (two-tailed). (**M**) Gross pulmonary metastases from human melanoma A375sm-Akaluc cells transfected with PHGDH sgRNA (sgRNA) in NSG mice by tail vein injection. C, empty vector. N= 10. Graphs show the mean ± SEM. The p-value is shown by an unpaired t-test (two-tailed). (**N**) Analyses of TCGA data show that overexpression of De novo serine synthesis (SSP) metabolism-related genes (PHGDH, NQO1, PSAT1, and SHMT2) is associated with a shorter time of survival in all tumor samples in TCGA data [p = 6.098e-4 by Logrank (Mantel-Cox) test].

To further confirm this notion, we tracked the metastases for 57 days after transplantation and found that the PHGDH-deficient group maintained lower luminescence signals, while the control group gradually increased its luminescence signals (Figure 6L). Consistently, the number of metastases in the PHGDH-deficient group remarkably decreased compared to the control group (Figure 6M). Our data demonstrate that the deficiency of PHGDH affects the survival and growth of metastatic tumors in vivo. Furthermore, we also found that overexpression of other genes such as SHMT2 and MTHFD2 or PSPH in one-carbon and SSP metabolism pathways could promote the tumor metastasis (Figure S3) and ATP production as well as the OCR in tumor cells (Figure S4), while the knockout of SHMT2 could inhibit the metastatic ability in highly metastatic cell lines (Figure S4). Moreover, the upregulation of PHGDH and SHMT2 correlated with melanoma progression and tumor metastasis in melanoma datasets (Figure S5), and overexpression of these four genes in the TCGA data sets could predict a worse overall survival rate and be associated with high metastasis rates in the datasets (Figure S5G). These findings suggest that, like one-carbon signaling pathways, the de novo serine synthesis (DSSP) metabolism pathway is also involved in metastatic regulation, and related genes are predictive markers and potential treatment targets for metastatic disease.

### Elimination or restriction of the one-carbon metabolism pathway blocks metastasis

We identified that one-carbon metabolism, de novo serine synthesis (DSSP) pathways, and related genes are associated with tumor metastasis, suggesting that they may represent vulnerabilities in metastatic cancer for the development of new therapeutic drugs. Previous studies reported that cancer cells not only require extracellular serine to support cell proliferation but also often increase serine synthesis from glucose and require de novo serine synthesis for one-carbon metabolism to facilitate amino acid transport, nucleotide synthesis, folate metabolism, and redox homeostasis in metastatic progression [25]. Increased glucose utilization is a characteristic of cancer cells that supports cell survival, proliferation, and metastasis [26]. Based on our findings of downregulated genes in the SSP pathway and one-carbon metabolism pathway, which are associated with the inhibition of metastases in at least melanoma, rhabdomyosarcoma and breast cancers (Figures 2A and S1), we first tested whether blocking glucose uptake by 2-DG affects tumor metastasis. As Figures 7A-F show, treatment with 2-DG not only inhibited metastasis of the mouse rhabdomyosarcoma RMS14 (Figure 7A) and melanoma B16F10 (Figure 7B) cells in syngeneic immunocompetent mice by tail vein injection but also blocked the metastasis of human melanoma A375sm (Figure 7C) and WM88 in immunodeficient NSG (Figures 7D and 7E) or nude mice (Figures 7D and 7E) by tail vein injection and inhibited the metastasis of human melanoma A375sm (Figure 7F) or reduced the IVIS luminescence intensity of lung (Figure 7G) by orthotopic footpad injection.

**Figure 7.**
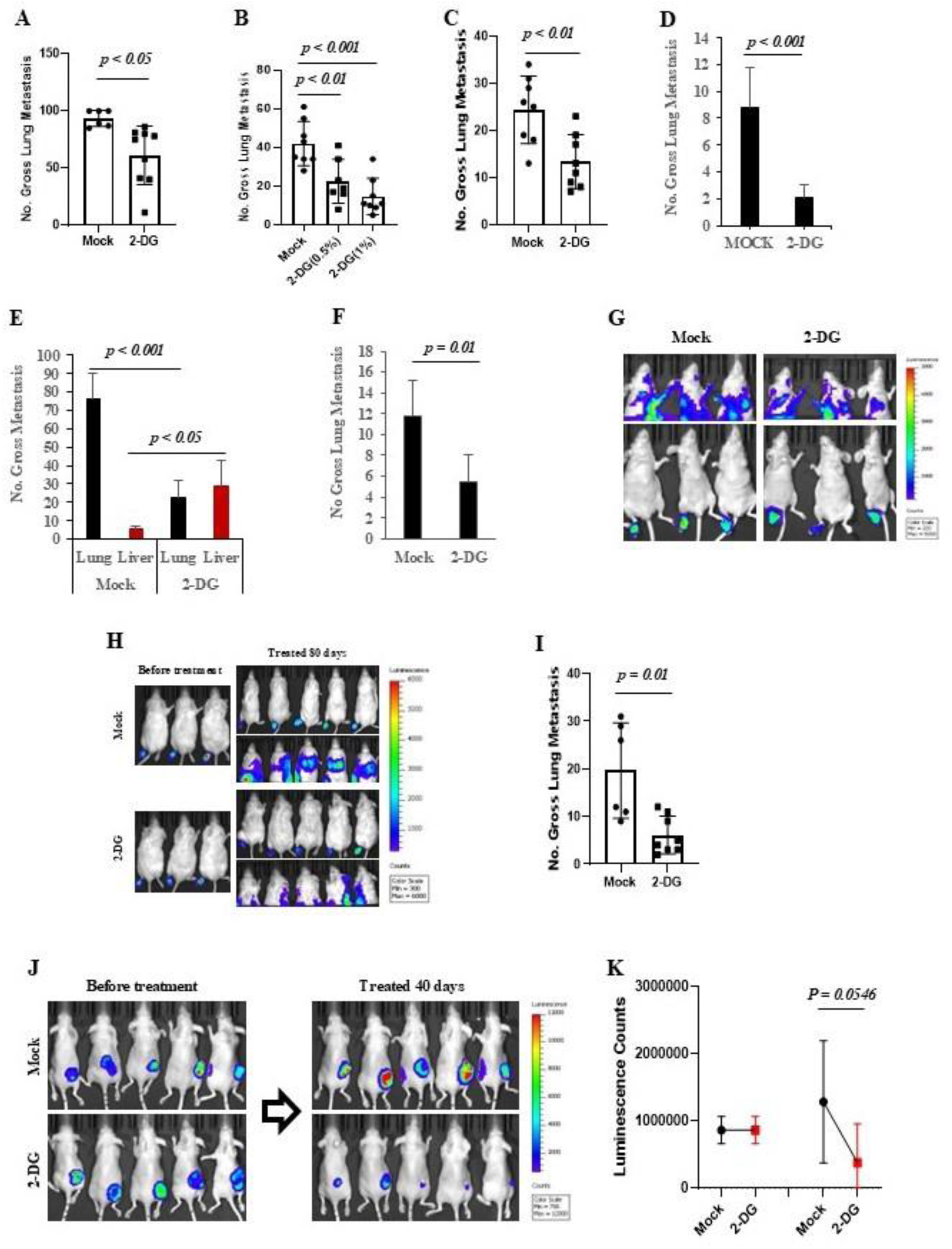
Blocking glucose uptake in the one-carbon metabolism pathway could inhibit tumor growth and metastasis. (**A**) Gross pulmonary metastases from rhabdomyosarcoma RMS14 cells in syngeneic FVB mice treated with 1% 2-Deoxyglucose (2-DG) (a glycolysis inhibitor). C. Mock, as a control; 2-DG, 1% 2-DG in drinker water for 4 weeks. n=10 mice, Graphs show the mean ± SEM. The p-value is shown by an unpaired t-test (two-tailed). (**B**) Gross pulmonary metastases from rmelanoma B16F10 cells in syngeneic BL6 mice treated with 0.5% or 1% 2-Deoxyglucose (2-DG) in drinking water. Mock, as a control; 2-DG (0.5%) or 2-DG (1%), 0.5% or 1% 2-DG in drinking water for 4 weeks. n = 10 mice. Graphs show the mean ± SEM. The p-value is shown by an unpaired t-test (two-tailed). (**C**) Gross pulmonary metastases from NSG mice transplanted with human melanoma A375sm by tail vein injection were treated with 1% 2-Deoxyglucose (2-DG). C. Mock, as a control; 2-DG, 1% 2-DG in drinking water for 4 weeks. n=10 mice, Graphs show the mean ± SEM. The p-value is shown by an unpaired t-test (two-tailed). (**D**) Gross pulmonary metastases from nude mice transplanted with human melanoma WM88 cells treated with 1% 2-Deoxyglucose (2-DG). C. Mock, as a control; 2-DG, 1% 2-DG in drinking water for 4 weeks. n=10 mice, Graphs show the mean ± SEM. The p-value is shown by an unpaired t-test (two-tailed). (**E**) Gross pulmonary metastases from NSG mice transplanted with WM88 cells treated with 1% 2-Deoxyglucose (2-DG). C. Mock, as a control; 2-DG, 1% 2-DG in drinking water for 4 weeks. n=10 mice, Graphs show the mean ± SEM. The p-value is shown by an unpaired t-test (two-tailed). (**F**) Gross pulmonary metastases from NSG mice transplanted with human melanoma A375sm cells by orthotopic footpad injection, treated with 1% 2-deoxyglucose (2-DG). C. Mock, as a control; 2-DG, 1% 2-DG in drinking water for 4 weeks. n=10 mice, Graphs show the mean ± SEM. The p-value is shown by an unpaired t-test (two-tailed). (**G**) Representation of bioluminescence images of the mice transplanted with A375sm-Akaluc cells by orthotopic feetpad injection to track the A375sm-Akaluc tumor cells treated with water control (Mock) and 1% 2-DG in drinking water (2-DG) using In Vivo Imaging System (IVIS). (**H**) Representation of bioluminescence images of mice transplanted with A375sm-Akaluc cells via orthotopic footpad injection to track the A375sm-Akaluc tumor cells before and after treatment with water control (Mock) and 1% 2-DG in drinking water (2-DG) for 80 days using an In Vivo Imaging System (IVIS). (**I**) Gross pulmonary metastases from mice transplanted with human melanoma A375sm-Akaluc cells by orthotopic feetpad injection were treated with 1% 2-DG for 80 days. C, empty vector. N=10. Graphs show the mean ± SEM. The p-value is shown by an unpaired t-test (two-tailed). (**J**) Representation of bioluminescence images of mice transplanted with A375sm-Akaluc cells via subcutaneous (SQ or Sub-Q) injection to track the growth of A375sm-Akaluc tumor cells before and after treatment with water control (Mock) and 1% 2-DG in drinking water (2-DG) for 40 days using an In Vivo Imaging System (IVIS). (**K**) The data showed the quantification of the bioluminescence signals of the local tumor in the mice transplanted with A375sm-Akaluc cells through subcutaneous (SQ or Sub-Q) injection to track the A375sm-Akaluc tumor cells’ growth before and after treatment with water control (Mock) and 1% 2-DG in drinking water (2-DG) for 40 days using In Vivo Imaging System (IVIS). N=10. Graphs show the mean ± SEM. The p-value is shown by an unpaired t-test (two-tailed).

To further confirm the inhibition of tumor metastasis by blocking glucose uptake through 2-DG, we tracked the effect of 2-DG on human melanoma A375sm metastasis in vivo after transplantation by orthotopic footpad injection. As Figures 7H and 7I show, before treatment with 2-DG, the treated (2-DG) and non-treated (Mock) groups exhibited similar signals at the footpad. However, after being treated for 80 days with 2-DG, the treated group had significantly reduced signals at the chest and local footpad (Figure 7H). The number of gross lung metastases was significantly reduced by treating with 2-DG (Figure 7I). Moreover, after A375smAkaluc cells were transplanted at a 100 cm³ size in NSG mice, treatment with 2-DG for 40 days significantly reduced tumor growth (Figure 7J and 7K). Notably, treatment with 2-DG downregulated the expression of ENOPH1, PHGDH, and SHMT2 in human melanoma (A375sm and WM88) and mouse melanoma (B16F10) cell lines as well as mouse rhabdomyosarcoma (RMS14) cell lines (Figure 8A). Our data suggest that blocking glucose uptake could inhibit one-carbon and SSP metabolism pathways gene expression, therefore blocking tumor proliferation and metastasis.

**Figure 8.**
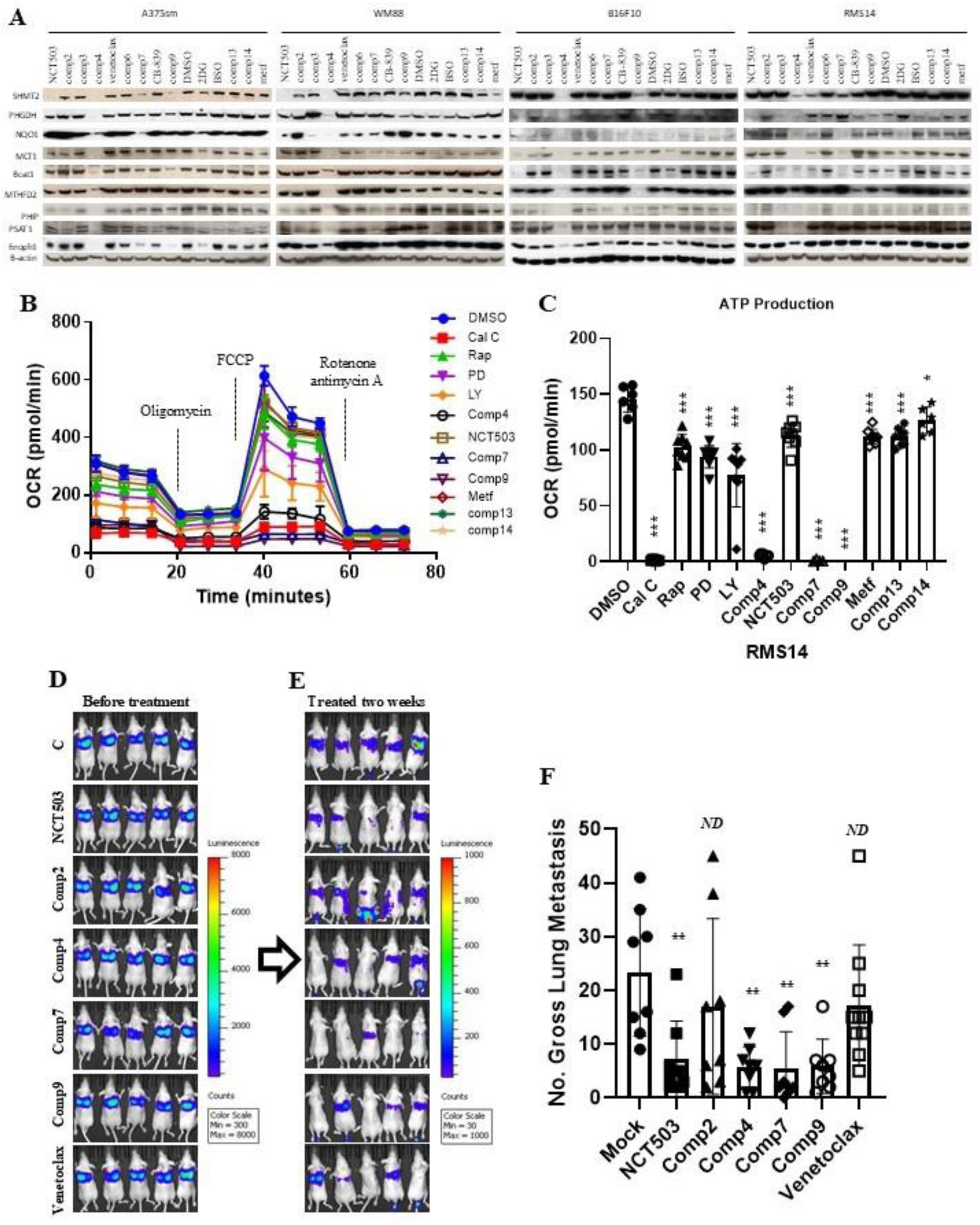
Elimination or restriction of the one-carbon metabolism pathway blocks metastasis. (**A**) Western blotting showed the expression of the indicated one-carbon and SSP metabolism pathway genes treated with different compounds, including identified new compounds comp4, comp7, and comp9. The cells were treated with 5 µM NCT503, 10 µM comp2, 10 µM comp3, 100 nM comp4, 10µM venetoclax, 10 µM comp6, 10 nM comp7, 30 nM CB-839, 10µM comp9, DMSO, 5 mM 2-DG, 1 mM BSO, 10 µM comp13, 10 µM comp14, 2 mM metf (Metformin) for 36 hours. (**B, C**) Seahorse analysis showed the effect of the different compounds on OCR (B) and ATP production (C). The cells were treated with 100 nM calphostin (Cal C), 10nM Rapamycin (Rap), 20µM MAPK inhibitor PD98059 (PD), 10µM PI3K/AKT inhibitor LY294002 (LY), 100 nM comp4, 5 µM NCT503, 10 nM comp7, 10µM comp9, 2 mM Metformin(Metf), 10 µM comp13, 10 µM comp14 and DMSO control. *ND*, no statistical difference; \**p* < 0.05; \*\**p* < 0.01; \*\*\**p* < 0.001. (**D, E**) Representation of bioluminescence signals of tumor metastasis from before treatment (D) and treated for two weeks with indicated compouds (E) after A375sm cell transplantation in NSG mice tracked by of the In Vivo Imaging System (IVIS). C, DMSO control; NCT503, 20 mg/Kg; Comp2, 25 mg/Kg; Comp4, 30 mg/Kg; comp7, 3 mg/Kg; comp9, 10 mg/Kg; venetoclax, 12 mg/Kg. (**F**) Gross pulmonary metastases from mice transplanted with cells by tail vein injection and treated with different compounds. N=10. Graphs show the mean ± SEM. The p-value is shown by an unpaired t-test (two-tailed). *ND*, no statistical difference; \**p* < 0.05; \*\**p* < 0.01; \*\*\**p* < 0.001.

To further discover more compounds that could target the one-carbon and SSP metabolism pathway genes, we screened the compounds library and identified three new compounds, comp4, comp7 and comp9, along with the well-known PHGDH inhibitor NCT503, that were able to inhibit all genes or at least three genes expression at protein level in human melanoma cell A375sm and WM88 and mouse melanoma B16F10 as well as mouse rhabdomyosarcoma RMS14 (Figure 8A). They also decreased OCR activation (Figure 8B) and ATP production (Figure 8C). Notably, they all could inhibit melanoma A375sm cell metastasis (Figures 8D, 8E and 8F), indicating that the discovered new compounds could inhibit the expression of genes in the one-carbon or SSP metabolism pathway and block tumor metastasis, and implying that targeting the one-carbon and SSP metabolism pathway could be a novel strategy for the treatment of metastatic disease.

## Discussion

Metastasis is a complex process that involves multiple steps, including shedding from the primary site, local invasion, intravasation into the bloodstream or lymphatics, spread through circulation, arresting at the capillary bed, extravasation into distant tissues, and localization and growth at a new site [3,7–10]. Tumor cells must pass through all steps to survive successfully and then become a metastatic tumor. Studies have shown that this metastatic cascade is a highly inefficient process, as most cancer cells do not survive; however, some undergo metabolic rewiring to optimize survival in specific microenvironments [11,27]. For example, cancer cells in primary sites are often in a hypoxic TME and use anaerobic glycolysis for cell growth and proliferation [11,28]. Cancer cells that can metastasize detach from the primary tumor site, experience high levels of oxidative stress, and undergo metabolic and transcriptional changes to survive in the harsh environment of the blood [11,29,30]. Once extravasated and seeded in distant organs, cancer cells need to reprogram their metabolism to survive and proliferate using the nutrients and oxygen available at secondary sites. The diversity of the metabolic pathways supporting cancer invasion limits the possibility of designing overarching treatment strategies. Common alterations in metabolism during metastasis may be vulnerabilities for developing treatments and strategies to combat deadly metastatic disease.

Applying metastatic function screening, we have identified that one-carbon and SSP metabolism pathways, as well as related genes, are associated with the inhibition of tumor metastasis. Overexpression of the genes (ENOPH1, PHGDH, SMTH2, and MTHFD2) in one-carbon and SSP metabolism pathways promoted tumor metastasis, while inhibiting these genes’ expression by CRISPR/Cas9 gene editing or small molecular inhibitors inhibited the tumor metastasis. Moreover, we identified three new compounds that are able to inhibit these gene expressions and block tumor metastasis in vivo. Our findings uncovered the mechanism by which tumor cells reprogram their metabolism to promote migration, invasion, and survival at distant sites, offering a rational strategy to guide clinical treatment. The identified novel molecular proteins and pathways represented a promising therapeutic target for metastatic disease.

Due to inter- or intra-tumor heterogeneity, various signaling pathways are required for metastasis in different cancers. We demonstrated that PKC and mTOR inhibitors generally inhibited metastasis, whereas the functions of the MAPK and PI3K pathways on metastasis depended more on the tumor type. Targeting the PI3/AKT signaling pathway inhibited tumor metastasis in melanoma, but inhibition of the MAPK pathway blocked the metastasis of rhabdomyosarcoma. Recent studies have shown that mTOR inhibition can inhibit tumor cell growth through epigenetic suppression of the serine glycine one-carbon metabolism pathway [31], glucose uptake [32], one-carbon metabolism requirements [33], and amino acid synthesis and metabolism [34,35]. Indeed, PKC, particularly PKCδ, is known to regulate tumor metastasis [36] and has been recently reported to promote NADPH production in oncogenic KRAS-mediated tumor progression [37]. These data support that inhibition of PKC and mTOR rewrites the tumor reprogramming, along with inhibition of MAPK in rhabdomyosarcoma and inhibition of PI3/AKT in melanoma to block the tumor metastasis. We found that the one-carbon metabolism pathway and serine synthesis (SSP) are essential for tumor metastasis in several cancers, and the notion was confirmed that blocking ENOPH1 and PHGDH (or SHMT2) was associated with inhibiting the tumor metastatic potential. Interestingly, in addition to the one-carbon and SSP pathways, we also found that folate cycle enzymes, such as MTHFD2, SHMT2, and MTHFD1, are associated with tumor metastasis, which are found to be upregulated in multiple cancers [38–43]. Our data suggest that targeting ENOPH1 or PHGDH, and their associated metabolic pathway, could be utilized for therapeutic purposes in treating metastatic disease.

Metastatic cancer cells evolve in harsh environments where most nutrients are scarce, a hallmark property that enables them to increase functions such as glycolysis or protein biosynthesis using alternative metabolic pathways [11,44]. For instance, one-carbon metabolism and the SPP pathway provide and support the production of methionine, serine, glycine, and nucleotide for DNA replication, NADPH for antioxidant defense, and methyl groups for modifying DNA, RNA, and protein [11,25,37,45,46], which all enable metastatic tumor cells to survive and migrate [47,48]. The production of SSP-derived α-ketoglutarate was shown to be responsible for enhanced breast cancer metastasis [49]. Numerous other studies have implicated one-carbon metabolism activity in increasing serine synthesis during the metastatic process [25,50]. Moreover, aggressive breast cancer cells also upregulate SSP one-carbon metabolism, and expression of SHMT2 is associated with poor survival in patients [51]. Indeed, enhanced SSP can promote metastasis without impacting primary tumor cell proliferation or growth [52–57]. Studies have reported that the methionine salvage pathway recycles one-carbon units lost during polyamine biosynthesis, thereby replenishing the methionine cycle and overcoming stress [44,58]. The methionine cycle regulates the rate of protein synthesis. Methionine restriction is a dietary regimen that protects against metabolic diseases and aging, represses cancer growth, and enhances the effectiveness of cancer therapy [59,60]. Methionine deprivation has also been found to decrease invasion and wound healing, suggesting that it may be beneficial in targeting metastatic progression [61–63]. Indeed, methionine deprivation or restriction also inhibits metastasis in diverse murine models, including sarcoma, triple-negative breast cancer, and melanoma [64,65]. We identified ENOPH1, a newly discovered enolase-phosphatase involved in methionine biosynthesis via the salvage pathway, which maintains methionine levels in vivo [66,67]. Deletion of ENOPH1 leads to growth restriction and severe stunting [68–71] is associated with metastatic potential. Inhibition of ENOPH1 reduces metastatic ability, while its overexpression promotes tumor metastasis. A recent study reported that ENOPH1 overexpression occurs in high-grade glioma, and its expression promotes cell proliferation and migration in glioma cells [72]. Additionally, ENOPH1 expression is upregulated in association with hepatocellular carcinoma metastatic potential [73]. Furthermore, our data suggest that targeting ENOPH1 and its pathway could be utilized for therapeutic purposes in treating metastatic disease.

In 1924, Otto Warburg reported that cancer cells metabolize glucose differently from normal cells [74–76]. Clinical studies have shown that most primary and metastatic cancers significantly increase glucose levels compared to normal tissues [77]. Overexpression of glucose transporters (GLUTs) such as GLUT1 or GLUT3 is associated with tumor metastasis [78,79]. Metastatic tumors often present high glucose consumption due to the upregulation of glycolysis (the Warburg effect) and downregulation of oxidative phosphorylation [80,81]. Activation of oxidative metabolism in TNBCs can reduce primary tumor growth and metastasis formation [47]. Targeting glycolytic metabolism represents an attractive strategy for treating cancers. For example, inhibiting glycolysis by 2DG induces a metabolic shift toward OXPHOS, decreases lactate production, and consequently inhibits cancer metastasis [82]. OXPHOS and increased mitochondrial metabolism have been reported to be required for the metastatic phenotype [83,84]. Moreover, simple sugar intake in drinks and fruit juice was associated with an increased risk of overall cancer incidence and mortality [85,86]. These suggest targeting glucose metabolism as an anticancer therapeutic approach because it alters the levels of many metabolite molecules, such as serine glycine [29,87,88].

Although many tumor cells depend on exogenous Serine, dietary Serine and glycine starvation can inhibit the growth of these cancers and extend survival in mice. However, under stress conditions, cancer cells, especially metastatic tumors, can produce serine and glycine from glucose by enhancing the expression of the de novo serine synthesis pathway (SSP) or activating oncogenes that drive enhanced serine synthesis and promote resistance to this starvation therapeutic approach [89]. Inhibiting the one-carbon metabolism enzyme SHMT2 affects tumor cell survival and metastasis [39]. Here, we demonstrate that inhibition of one-carbon metabolism and SSP pathway genes is associated with a reduction in the metastatic potential of cancer. Importantly, we confirmed that the inhibition of PHGDH, the first step in the SSP, blocked tumor growth, metastatic tumor survival, and metastatic potential.

The importance of serine as a precursor to diverse pathways that control biosynthesis, epigenetic regulation, redox homeostasis, ATP generation, and TCA cycle regulation explains why cancer cells have a high demand for either endogenous or exogenous serine [23,66,88,90]. Consequently, genetic alterations of the SSP and one-carbon metabolism genes are frequently observed in cancer. For example, gene amplification or overexpression of PHGDH and PSPH are found in several cancer types, including estrogen receptor (ER)-negative breast cancers, melanoma, lung adenocarcinoma, T cell ALL, and osteosarcoma [67,69–75]. In many cases, increased expression of SSP pathway-related genes correlates with a poor prognosis [70,72,76,77], a finding consistent with our identification in TCGA data (Figure 6N).

In summary, our findings reveal a mechanism by which tumor cells reprogram their metabolism to enhance migration, invasion, and survival at distant sites during metastasis. This research offers a rational strategy for guiding clinical treatment. The novel molecular proteins and pathways identified in our study represent promising therapeutic targets for metastatic diseases.

## Materials and Methods

### Plasmids, antibodies, cell lines and culture. Plasmids

ENOPH1, shRNA for ENOPH1, ENOPH1 TALEN, ENOPH1 CRISPR/CAS9 sgRNA, PHGDH wt, Akaluc plasmid, PHGDH dominant active form, PHGDH CRISPR/CAS9 sgRNA, SHMT2, SHMT2-CD, SHMT2 CRISPR/CAS9 sgRNA, MTHFD2, and PSPH expressing plasmids were provided by Addgene (Cambridge, MA). The ENOPH1 shRNA-expressing plasmid was purchased from Open Biosystems against nucleotides of the ENOPH1 coding region (mouse BC021445, 655-675 and human BC005821, 1283-1303). Antibodies: anti-SHMT2, anti-PHGDH, anti-BCAT1, anti-slc16a1, anti-MTHFD2, anti-PSAT1, anti-PSPH, anti-ENOPH1, anti-PHIP, and anti-NQO1 antibodies were obtained from CASCADE Biosciences (Winchester, MA), while anti-β-actin and anti-GAPDH antibodies were obtained from Cell Signaling (Danvers, MA, USA). Anti-β-actin antibody is from Santa Cruz (Dallas, TX). The B16F1 and B16F10 cell lines were obtained from the American Type Culture Collection (ATCC, Manassas, VA). Panel cell lines of A375p and A375sm were generously provided by Dr. Isaiah Fidler (M.D. Anderson Medical Center, Houston, TX). The WM88 cell line was a gift from Dr. Meenhard Herlyn (The Wistar Institute, Philadelphia, PA). Stable expressing cells were established through transfection using Lipofectamine 2000 reagent (Invitrogen, Carlsbad, CA) and selected with antibiotics G418 or puromycin (Sigma, St. Louis, MO).

### Small molecular compounds

Calphostin C, Rho Inhibitor I, Rapamycin, PKA inhibitor H-89, p38 inhibitor SB203580, HADC inhibitor Trichostatin A and MS-275, MAPK inhibitor PD98059, PI3K/AKT inhibitor LY294002, Rottlerin, bisindoylmaleimide I (BMI), PKC β inhibitor I, and FDA-approved-Drug-Library (1562 compounds, HY-L022) were obtained from MedChemExpress (Monmouth Junction, NJ). All compounds were dissolved in DMSO. The cells were treated with 100 nM Calphostin C, 12nM Rho Inhibitor I, 10 nM Rapamycin, 50 nM PKA inhibitor H-89, 10 µM p38 inhibitor SB203580, HADC inhibitor 300 nM Trichostatin A and 10 µM MS-275, 20 µM MAPK inhibitor PD98059, 10µM PI3K/AKT inhibitor LY294002, 5µM Rottlerin, 20nM bisindoylmaleimide I (BMI), 20nM PKC β inhibitor I, and 10µM FDA-approved-Drug-Library for 24 or 48 hours. DMSO was used as a control.

### Microarray analysis

microarray analysis was as described in previous studies [19,91]. RNAs from cells, with triplicate biological replications of each microarray experiment, were isolated using Trizol. Total RNA was further purified on an RNAeasy column (Qiagen), and the RNA quality was checked by an Agilent Bioanalyzer (Agilent Technologies, Palo Alto, CA, USA). Target labeling and hybridization to GeneChips were carried out in the NCI Frederick Microarray Core facility using the GeneChip Mouse 430_2 Array purchased from Affimetrix. The microarray signals were normalized using the RMA algorithm. The significantly expressed genes were selected based on ANOVA analysis by Partek Genomics Suite software (Partek, St. Charles, MO, USA). The ANOVA gene list, filtered by a P value of 0.05 and an absolute value of fold change of 1.5, was used in the gene ontology analysis via the commercial gene pathway analysis web tool (http://trials.genego.com/cgi/index.cgi).

### Analyses of clinical outcome data

Clinical follow-up and gene expression data sets were obtained from publicly available datasets (GDS3966, GDS1375, GSE3189, and GSE7553) in the Gene Expression Omnibus (GEO).

### Gene Set Enrichment Analysis (GSEA)

The GSEA was performed using the desktop software GSEA from the Broad Institute.

### GO gene pathway analysis

GO gene pathway analysis uses the Gene Ontology (GO) database to identify over-represented biological pathways and functions within a given gene list from RNA sequencing experiment.

### Transcription Activator-Like Effector Nucleases (TALENs) genomic editing

The TALEN genomic editing was performed as described in the protocol provided by the manufacturer using the TALEN Assembly Kit [92].

### CRISPR/CAS9 knockout

CRISPR/CAS9 knockout was performed as described in previous studies [36,91]. ENOPH1, PHGDH, and SHMT2 knockout sgRNAs CRISPR were used. We used the CRISPR/Cas9 technology with a homology-directed repair (HDR) system to specifically knock out ENOPH1, PHGDH, and SHMT2 in cells: sgRNA #1 ACCGCCAAATTTAATTGCAG, #2 CCTACCTCTGCAATTAAATT, and #3 TTATCCAAACATTATTGCTA.

### Western blot analysis

Immunoblots were performed on lysates generated from cultured cells and tissues solubilized in RIPA buffer [93,94].

### Cell proliferation

The CCK8 kit was used for the measurement of cell growth [6,95,96].

### Experimental and spontaneous metastasis assays

Cells were intravenously injected via a tail vein or footpad into 4 – 6-week-old male mice. There were six different parent cell lines: B16F1 cells were injected at 5×105 into FVB/BL6 or 1×105 into athymic nude; 37-7 cells were injected at 2×105 into FVB or 1×105 into athymic nude; human melanoma A375 panel cells were injected via tail vein or footpad into NSG mice at 1×106 [36,91,94,97]. Tumor numbers were obtained by visual inspection of tissues in mice euthanized 21 days post-transplantation, and micrometastases were counted by pathologists’ evaluation after dissection of the lung. All mouse procedures were performed in accordance with the National Institutes of Health guidelines. The animal studies were under the animal proposals LMB042 and LCBG023, approved by the National Cancer Institute-Bethesda Animal Care and Use Committee (ACUC) in the United States of America.

### Metabolon analysis

Metabolon was completed the mView Study Proposal to detect the differences in metabolism level in tumor cells (8 groups, total 32 samples), including: 1. Standard mView deliverables; 2. Unknown compounds; 3. Compound plexing (ref).

### Seahorse analysis

Adherent cells were seeded at 2 × 104 cells/well in normal growth media (cell line specific) in a Seahorse XF96 Cell Culture Microplate. To achieve an even distribution of cells within wells, plates were rocked at 25 °C for 20–40 min. For each staining group, one extra well on the outer perimeter of the plate was seeded to calibrate image acquisition parameters. The plate was then incubated at 37 °C overnight to allow the cells to adhere. The following day, growth media were exchanged with Seahorse Phenol Red-free DMEM, and basal OCR was measured or an XF Cell Mito stress test (Agilent) was performed according to the manufacturer’s instructions. In both cases, the last injection port was used for the injection of cell stains or dyes. Upon completion of the Seahorse assay, the cells were washed three times with pre-warmed phenol-free DMEM medium (without FBS) and then transferred to a Cytation 5 for imaging. For cells that grow in suspension, 105 cells/well were added to a poly-lysine-coated Seahorse XF96 Cell Culture Microplate in Phenol Red-free DMEM-based assay media. The plate was then centrifuged at 300 × g for 30 min with gentle acceleration and deceleration. The plate is then rotated 180° and the centrifugation is repeated. Immediately following completion of the centrifugation, OCR was measured, or an XF Cell Mito stress test was performed, followed by imaging in the Cytation 5 as above.

### Immunohistochemistry

Lung tissues were fixed in 10% buffered formalin solution (pH 7.2) for 16 h, and/or frozen in OCT compound and serially sectioned to 15 µm at 20 oC. Immunohistochemistry was performed as described [36,91,98]. Immunoreactivity scores were analyzed using ImageScope V 10.0 software from Aperio Technologies (Vista, CA). The size of the metastases was quantified using ImageJ software and analyzed with Statgraphics software.

### In Vivo Imaging System (IVIS)

Mice are injected by an intraperitoneal route with a Luciferin solution (15 mg/mL or 30 mg/kg, in PBS, dose of 150 mg/kg) that is allowed to distribute in awake animals for about 5-15 minutes. The mice are placed into a clear Plexiglas anesthesia box (1.5-2.5% isofluorane) that allows unimpeded visual monitoring of the animals, e.g., one can easily determine if the animals are breathing. The tube that supplies the anesthesia to the box is split so that the same concentration of anesthesia is plumbed to the anesthesia manifold located inside the imaging chamber. After the mice are fully anesthetized, they are transferred from the box to the nose cones attached to the manifold in the imaging chamber, the door is closed, and the “Acquire” button (part of the Living Image program) on the computer screen is activated. The imaging time is between one and five minutes per side (dorsal/ventral), depending on the experiment. When the mice are turned from dorsal to ventral (or vice versa), they can be visibly observed for any signs of distress or changes in vitality. The mice are again imaged (maximum five minutes), and the procedure is complete.

The mice are returned to their cages, where they wake up quickly. We plan to image the mice from the start day of treatment, then once every week, on the day the experiment ended. Also, to determine rates of tumor proliferation and metabolism in vivo, we plan to trace the metabolic network for a tumor in vivo using 13C6-glucose in the liquid diet as described in Nature Communications, 2017, 8:1646. 13C-glucose labeling of mice to determine rates of tumor proliferation and metabolism in vivo: Using 13C6-glucose in the liquid diet [the liquid diet base containing casein, L-cystine, soy oil, cellulose, mineral mix (AIN-93GMX), calcium phosphate, vitamin mix (AIN-93-VX), Choline Bitartrate, tert-butylhydroquinine (TBHQ), and Xanthan gum and 13C6-glucose], to determine rates of tumor proliferation and metabolism in vivo. The procedure: 1) Pre-feeding of an unlabeled liquid diet served to accustom the mice to the liquid diet feeding two days before the tracer study. The unlabeled glucose liquid diet and water are added to the diet base two days before the tracer study to give a final diet of 0.167 g glucose/g diet and a net protein content of 53 mg/g diet to provide sufficient carbon and nitrogen. Twenty g mice are fed a 13.6 g liquid diet (at 680 g diet/kg mouse). 2) Tracer study: on the third day, 13C6-glucose at 0.173 g/g diet replaced the unlabeled glucose in the diet for each mouse, and the mice were allowed to consume the diet ad libitum for 18 h. 3) At the end of the feeding period, mice are sacrificed by cervical dislocation, and organs are harvested and snap frozen in liquid nitrogen. The tracer study will be performed at the end of the in vivo experiment.

### Tissue microarray (TMA) analysis

Human ME207, ME208, and ME1004b tissue microarrays (TMA) contain nevus, primary melanomas, and lymph metastatic melanomas (US Biomax, Rockville, MD). TMA slides were stained with anti-PTEN, anti-Endpd5, anti-IGF1R, and ATF6 antibodies, and the substrate was DAB, AEC, or permanent red (DAKO Cytomation, Carpinteria, CA). TMA slides were scanned by ImageScope and reviewed by three pathologists. Immunoreactivity scores were analyzed by three pathologists using ImageScope V 10.0 software (Aperio Technologies, Vista, CA). Immunostaining intensities were scored as strong (3), moderate (2), weak (1), and negative (0). The frequency of labeled tumor cells was scored as follows: 100% as 5, 66-99% as 4, 33-65% as 3, 10-32% as 2, <10% as 1, and no cells as 0. IHC scores equal intensity plus frequency [91].

### Xenograft model

A375sm cells stably expressing indicated genes as well as empty vector were subcutaneously injected into athymic nude mice or NSG mice. Tumors were measured from day 7 after injection. The volumes were estimated using the formula: volume = width2 × length/2 [95,96].

### Statistics

Statistical analyses were performed as follows: unpaired t-test (two-tailed) for all column data sets or Pearson r (two-tailed) analysis for correlation or Log-rank (Mantel-Cox) test for Kaplan Meier survival analysis using GraphPad Prism 6 software. The p values of less than 0.05 were considered statistically significant.

## Acknowledgments

This work was supported by funding from the NIH intramural research program.

