## Supplemental data for "Targeting one-carbon metabolic vulnerabilities of metastasis with therapeutic potential"

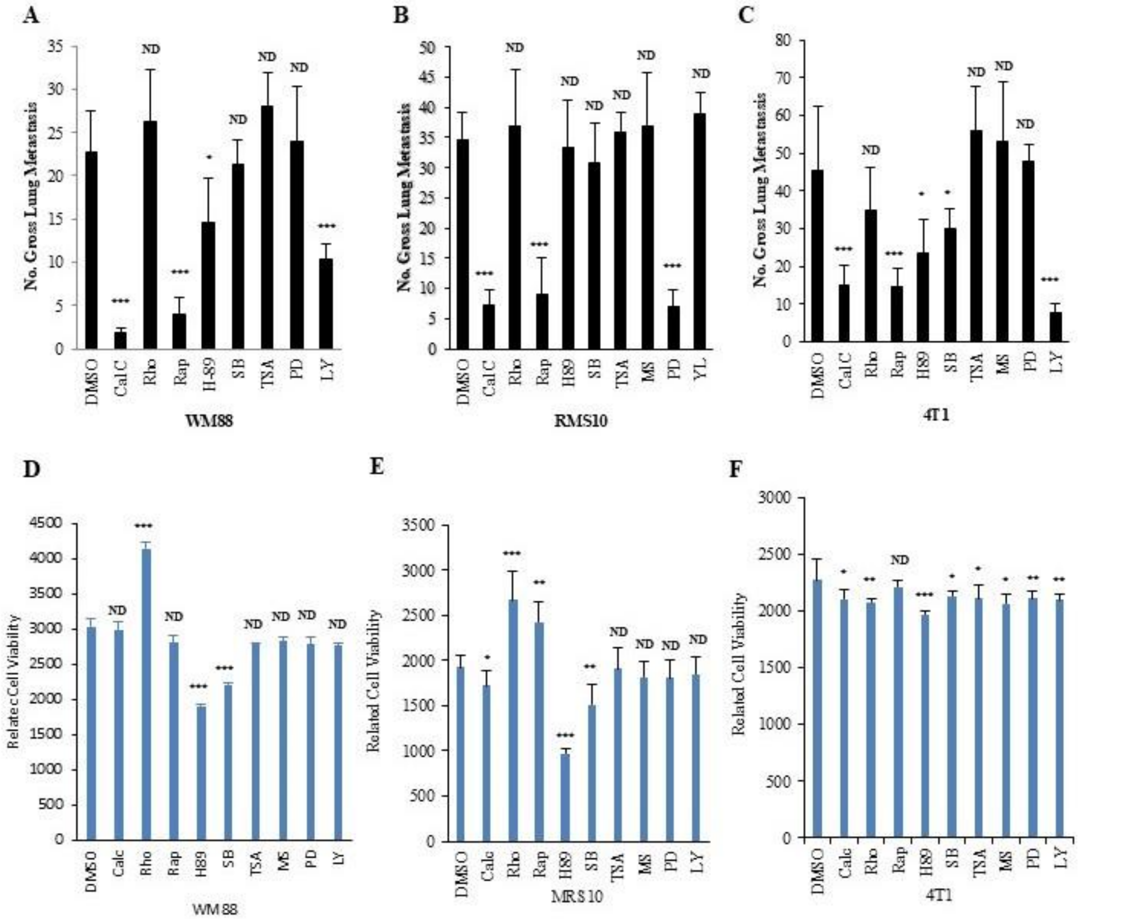


**Figure S1. Metastasis within and between tumors presented diverse responses to treatments.** (**A, B, C**) Gross pulmonary metastases from human melanoma WM88 (A), mouse rhabdomyosarcoma RMS10 (B), and mouse breast cancer 4T1 (C) in response to molecular inhibitor in the experimental metastasis assay by tail vein injection. Data are represented as mean ± SEM. The p-values were presented from an unpaired t-test analysis (two-tailed) compared with the control (DMSO). Cal C, 100 nM calphostin C; Rho, 12nM Rho Inhibitor I; Rap, 10nM Rapamycin; H-89, 50nM PKA inhibitor; SB, 10µM p38 inhibitor SB203580; TSA, 300nM HDAC inhibitor Trichostatin A; MS, 10µM HDAC inhibitor MS-275; PD, 20µM MAPK inhibitor PD98059; LY, 10µM PI3K/AKT inhibitor LY294002. *ND*, no statistical difference; *$p < 0.05$; **$p < 0.01$; ***$p < 0.001$. n = 10. (**D, E, and F**) The related cell viability of human melanoma WM88 (D), mouse rhabdomyosarcoma RMS10 (E), and mouse breast cancer 4T1 (F) cells, pretreated with molecular inhibitors. Data are represented as mean ± SEM of three independent experiments. The p-values were presented from an unpaired t-test analysis (two-tailed) compared with the control (DMSO). *ND*, no statistical difference; *$p < 0.05$; **$p < 0.01$; ***$p < 0.001$.

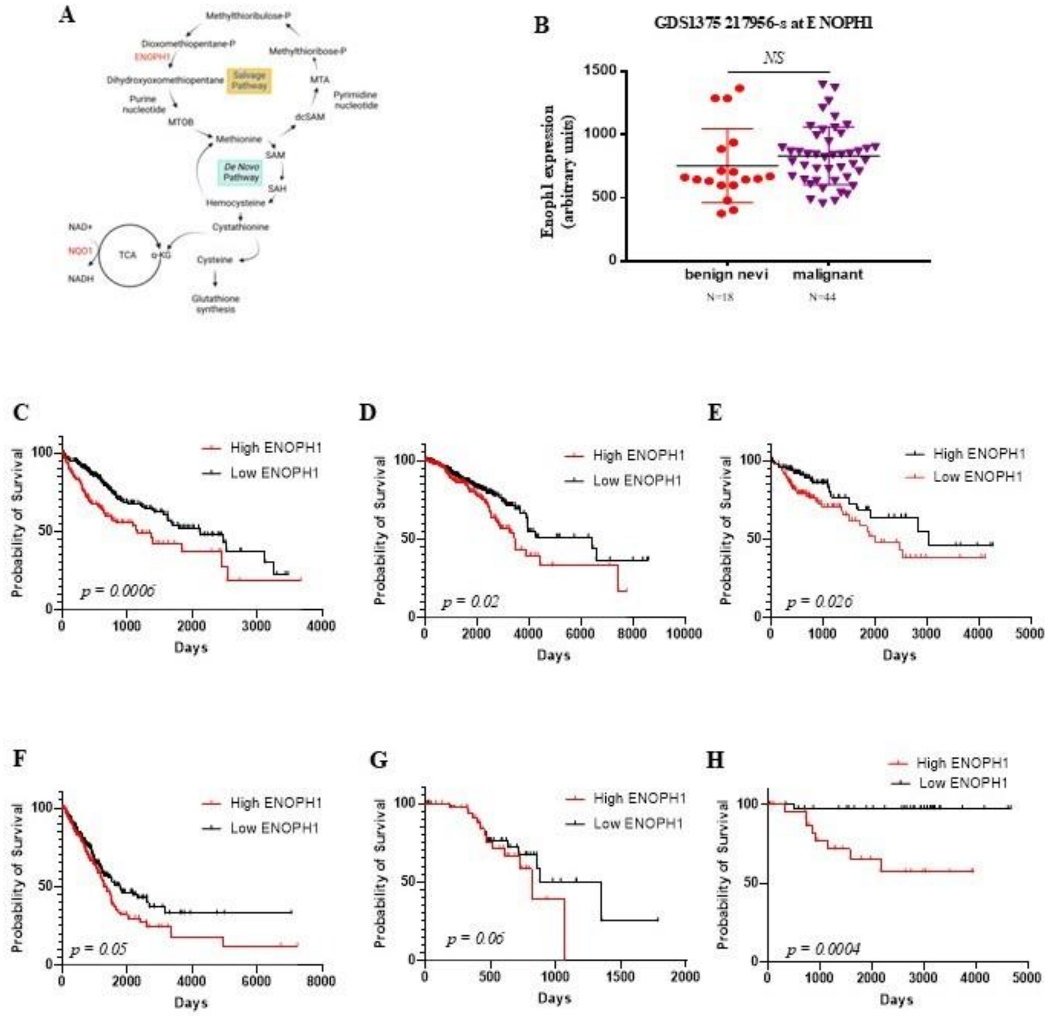


**Figure S2. One carbon metabolism-related gene, ENOPH1, is associated with the potential for metastasis.** (**A**) Schematic of the methionine salvage pathways. (**B**) ENOPH1 is expressed in benign nevi and melanoma in the GDS1375 dataset. Graphs show the mean ± SEM. The p-value is shown by an unpaired t-test (two-tailed). (**C to H**) The high level of ENOPH1 expression correlates with worse survival in TCGA datasets. The patients with high level of ENOPH1 (red line) had significantly worse survival than the patients with low level of ENOPH1 (black line) in various cancers (TCGA data): C, Liver cancers (Log-rank test p-value: 0.0006, high ENOPH1=145, low ENOPH1=217); D, Breast cancers (Log-rank test p-value: 0.02, high ENOPH1=399, low ENOPH1=622); E, Colon cancers (Log-rank test p-value: 0.026, high ENOPH1=109, low ENOPH1=145); F, Lung cancers (Log-rank test p-value: 0.05, high ENOPH1=207, low ENOPH1=290); G, melanoma (Log-rank test p-value: 0.06, high ENOPH1=39, low ENOPH1=59); H, Renal cancers (Log-rank test p-value: 0.0004, high ENOPH1=25, low ENOPH1=39).

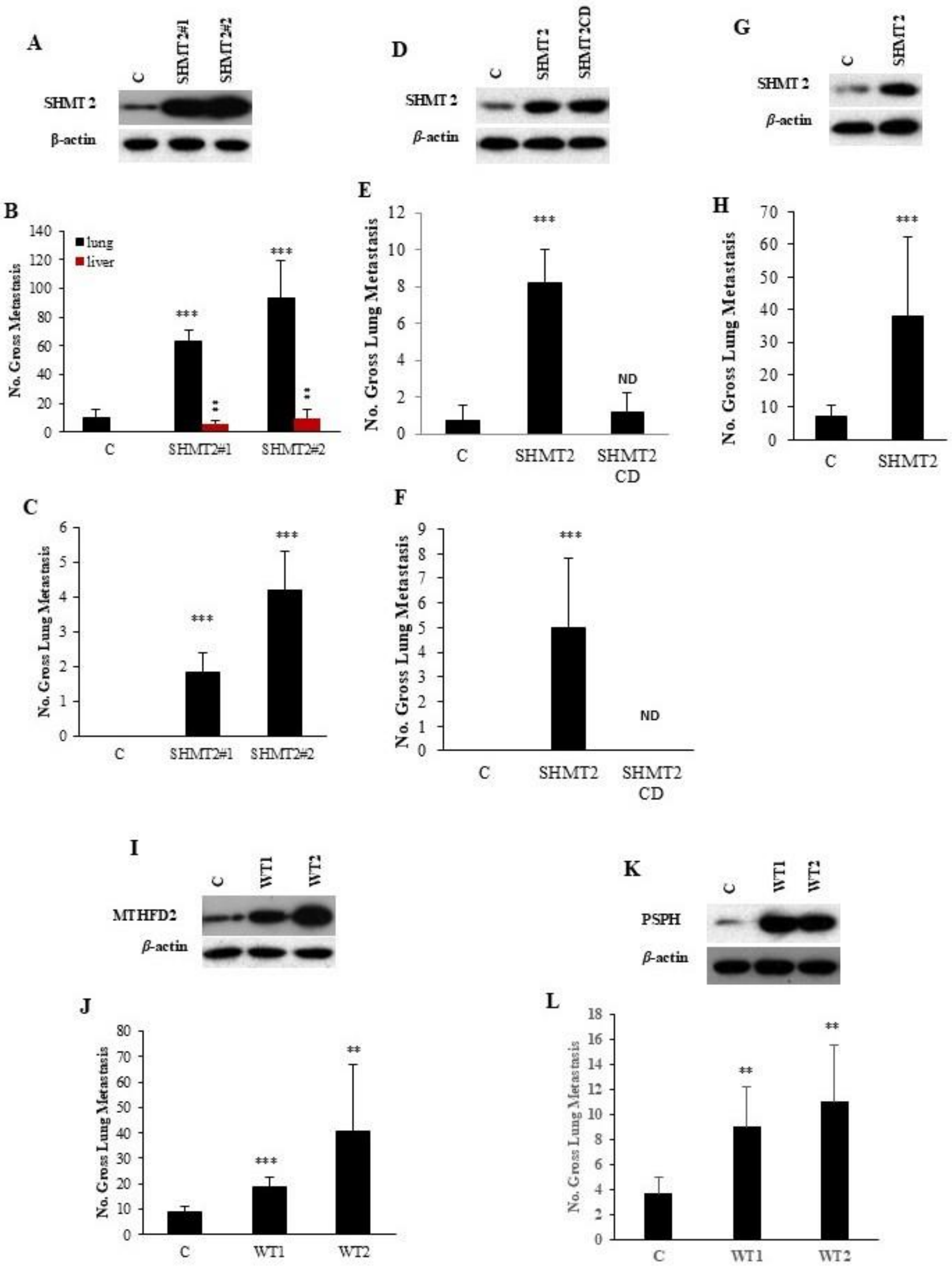


**Figure S3. One carbon metabolism-related genes are associated with the potential for metastasis**. (**A**) Western blotting showed overexpression of SHMT2 (a rate-limiting enzyme in the glycine biosynthesis pathway) in poorly metastatic human melanoma A375p cells. Protein levels of SHMT2 in A375p cells transfected with the wildtype SHMT2 gene (SHMT2#1 and SHMT2#2 C, empty vector; SHMT2#1 and SHMT2#2, two different clones with the SHMT2 gene. The protein level of β-actin is the loading control. (**B**) Gross metastases from human melanoma A375p cells transfected with the wildtype SHMT2 gene (SHMT2#1 and SHMT2#2) were induced by tail vein injection: c, empty vector; SHMT2#1 and SHMT2#2, two different clones with wildtype. N=10, Graphs show the mean ± SEM. The p-value is shown by an unpaired t-test (two-tailed). (**C**) ) Gross pulmonary metastases from human melanoma A375p cells transfected with wildtype SHMT2 gene (SHMT2#1 and SHMT2#2) by orthotopic footpad injection. C, empty vector; SHMT2#1 and SHMT2#2, two different clones with SHMT2 wildtype. N =10. Graphs show the mean ± SEM. The p-value is shown by an unpaired t-test (two-tailed). *ND*, no statistical difference; *$p < 0.05$; **$p < 0.01$; ***$p < 0.001$. (**D**) Western blotting showed overexpression of SHMT2 (a rate-limiting enzyme in the glycine biosynthesis pathway) in poorly metastatic mouse melanoma B16F1 cells. Protein levels of SHMT2 in B16F1 cells transfected with the wildtype SHMT2 gene (SHMT2) and kinase dead mutant form (SHMT2CD). C, empty vector; SHMT2, the clone with the SHMT2 gene; SHMT2CD, the clone with the SHMT2 kinase dead form. The protein level of β-actin is the loading control. (**E**) Gross metastases from mouse melanoma B16F1 cells transfected with the wildtype SHMT2 gene (SHMT2) or kinase dead mutant form (SHMT2CD) were induced by tail vein injection. C, empty vector; SHMT2, the clone with the SHMT2 gene; SHMT2CD, the clone with the SHMT2 kinase dead form. The protein level of β-actin is the loading control. (**F**) Gross metastases from mouse melanoma B16F1 cells transfected with the wildtype SHMT2 gene (SHMT2) or kinase dead mutant form (SHMT2CD) by orthotopic footpad injection. C, empty vector; SHMT2, the clone with the SHMT2 gene; SHMT2CD, the clone with the SHMT2 kinase dead form. The protein level of β-actin is the loading control. N =10. Graphs show the mean ± SEM. The p-value is shown by an unpaired t-test (two-tailed). *ND*, no statistical difference; *$p < 0.05$; **$p < 0.01$; ***$p < 0.001$. (**G**) Western blotting showed overexpression of SHMT2 (a rate-limiting enzyme in the glycine biosynthesis pathway) in poorly metastatic mouse rhabdomyosarcoma RMS119 cells. Protein levels of SHMT2 in RMS119 cells transfected with the wildtype SHMT2 gene (SHMT2). C, empty vector; SHMT, the clone with the SHMT2 gene. The protein level of β-actin is the loading control. (**H**) Gross metastases from mouse rhabdomyosarcoma RMS119 cells transfected with the wildtype SHMT2 gene (SHMT2) were induced by tail vein injection. C, empty vector; SHMT2, the clone with the SHMT2 gene. N=10, Graphs show the mean ± SEM. The p-value is shown by an unpaired t-test (two-tailed). *ND*, no statistical difference; *$p < 0.05$; **$p < 0.01$; ***$p < 0.001$. (**I**) Western blotting showed overexpression of MTHFD2 in poorly metastatic human melanoma A375p cells. Protein levels of MTHFD2 in A375p cells transfected with the wildtype MTHFD2 gene (WT1 and WT2). C, empty vector; WT1 and WT2, the clone with the MTHFD2 gene. The protein level of β-actin is the loading control. (**J**) Gross metastases from human melanoma A375p cells transfected with the wildtype MTHFD2 gene (WT1 and WT2) were induced by tail vein injection. C, empty vector; WT1 and WT2, the clones with MTHFD2. N=10, Graphs show the mean ± SEM. The p-value is shown by an unpaired t-test (two-tailed). *ND*, no statistical difference; *$p < 0.05$; **$p < 0.01$; ***$p < 0.001$. (**K**) Western blotting revealed the overexpression of PSPH in poorly metastatic human melanoma A375p cells. Protein levels of PSPH in A375p cells transfected with the wildtype PSPH gene (WT1 and WT2). C, empty vector; WT1 and WT2, the clone with the PSPH gene. The protein level of β-actin is the loading control. (**L**) Gross metastases from human melanoma A375p cells transfected with the wildtype PSPH gene (WT1 and WT2) were induced by tail vein injection. C, empty vector; WT1 and WT2, the clones with PSPH. N=10, Graphs show the mean ± SEM. The p-value is shown by an unpaired t-test (two-tailed). *ND*, no statistical difference; *$p < 0.05$; **$p < 0.01$; ***$p < 0.001$.

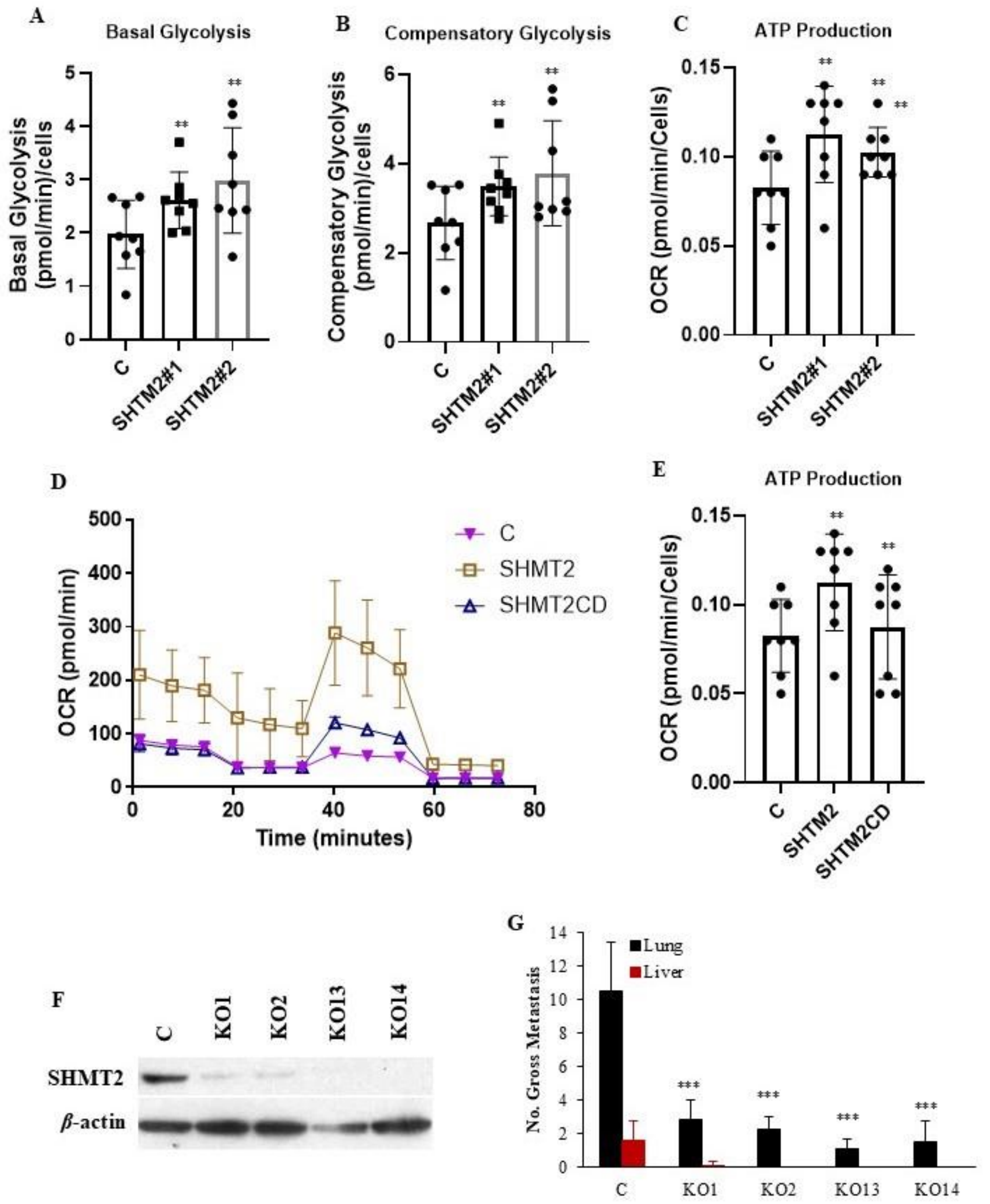


**Figure S4. The alteration of SHMT2 expression affects the tumor cell metabolism, survival in vivo, and the potential of metastasis**. (**A, B, C**) Seahorse analysis confirmed that overexpression of SHTM2 in A375p cells enhances both basal (A) and compensatory glycolysis (B), ATP production (C). C, empty vector control. The data represent three independent experiments. Graphs show the mean ± SEM. The p-value is shown by an unpaired t-test (two-tailed). *ND*, no statistical difference; $*p < 0.05$; $**p < 0.01$; $***p < 0.001$. (**D and E**) Seahorse analysis confirmed that SHTM2 in B16F1 cells enhances the OCR (D) and ATP production (E), while SHTM2 kinase dead does not affect either the OCR (D) or ATP production (E). C, empty vector control. The data represent three independent experiments. Graphs show the mean ± SEM. The p-value is shown by an unpaired t-test (two-tailed). *ND*, no statistical difference; $*p < 0.05$; $**p < 0.01$; $***p < 0.001$. (**F**) Western blotting showed the knockout of SHMT2 by CRISPR/CAS9 sgRNA gene editing in highly metastatic human melanoma A375sm cells. Protein levels of SHMT2 in A375sm cells transfected with CRISPR/CAS9 plasmids (sgRNAs): c, empty vector. The protein level of β-actin is the loading control. (**G**) Gross pulmonary metastases from human melanoma A375sm cells transfected with SHMT2 sgRNA (sgRNAs) in NSG mice by tail vein injection: c, empty vector. N=10, Graphs show the mean ± SEM. The p-value is shown by an unpaired t-test (two-tailed). *ND*, no statistical difference; $*p < 0.05$; $**p < 0.01$; $***p < 0.001$.

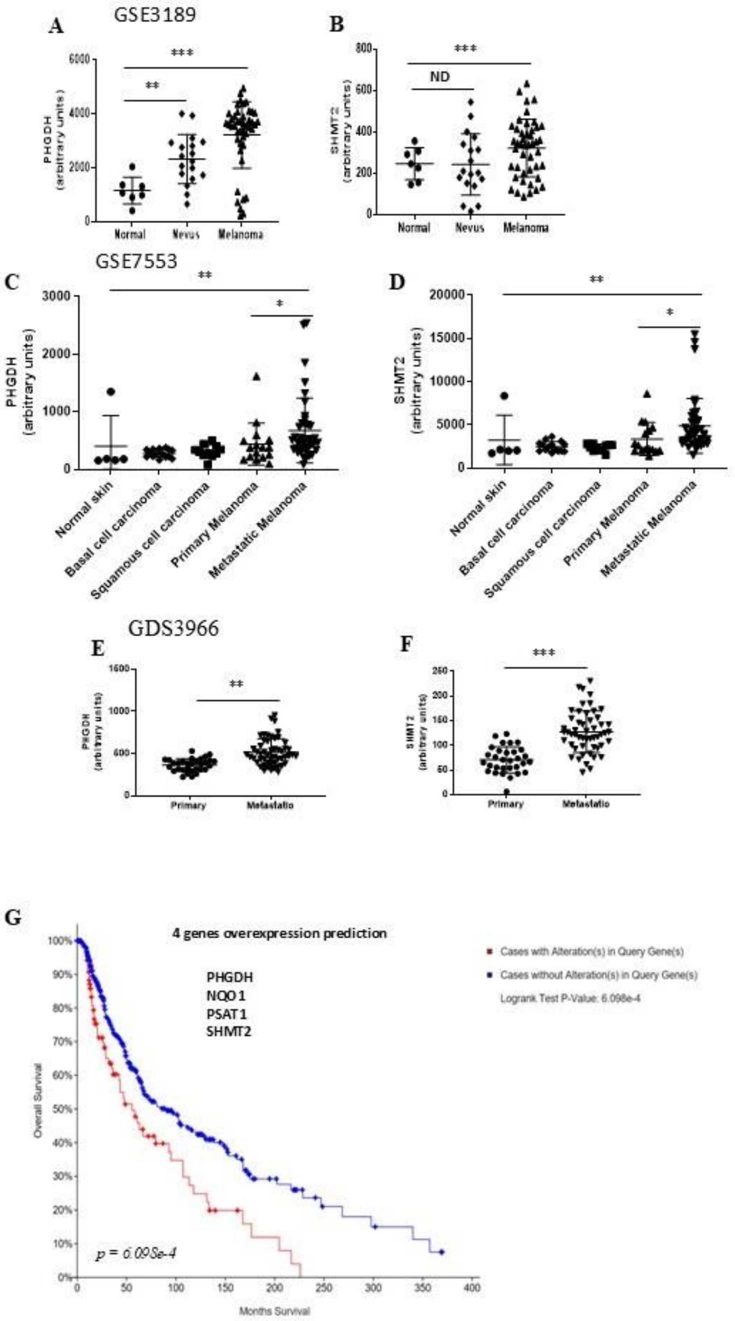


**Figure S5. One carbon metabolism-related gene is associated with tumor progression.** (**A and B**) PHGDH (A) or SHMT2 (B) is highly significantly expressed in melanoma compared to normal or nevus in the GSE3189 dataset. Graphs show the mean

± SEM. The p-value is shown by an unpaired t-test (two-tailed). *ND*, no statistical difference; *$p < 0.05$; **$p < 0.01$; ***$p < 0.001$. (**C and D**) PHGDH (C) or SHMT2 (D) is highly significantly expressed in metastatic melanoma compared to primary melanoma in the GSE7553 dataset. Graphs show the mean ± SEM. The p-value is shown by an unpaired t-test (two-tailed). *ND*, no statistical difference; *$p < 0.05$; **$p < 0.01$; ***$p < 0.001$. (**E and F**) PHGDH (E) or SHMT2 (F) is highly significantly expressed in metastatic melanoma compared to primary melanoma in the GDS3966 dataset. Graphs show the mean ± SEM. The p-value is shown by an unpaired t-test (two-tailed). *ND*, no statistical difference; *$p < 0.05$; **$p < 0.01$; ***$p < 0.001$. (**G**) Analyses of TCGA data show that overexpression of De novo serine synthesis (SSP) metabolism-related genes (PHGDH, NQO1, PSAT1, and SHMT2) is associated with a shorter time of survival in all tumor samples in TCGA data [p = 6.098e-4 by Logrank (Mantel-Cox) test].
